# Uncovering directionally and temporally variable genes with STAVAG

**DOI:** 10.1101/2025.09.02.673732

**Authors:** Qunlun Shen, Kuo Gai, Shang Li, Fangkun Yang, Hanbin Cui, Shuqin Zhang, Shihua Zhang

## Abstract

Spatially variable genes (SVGs) are crucial for understanding spatial heterogeneity in spatial transcriptomics. Yet, SVGs are challenging to align with established biological axes and lack straightforward mechanistic interpretation, thereby limiting their ability to inform downstream experiments or clinical applications. Here, we introduce the concept of directionally variable genes (DVGs) and temporally variable genes (TVGs). Also, we propose a unified framework, STAVAG, that models spatial or temporal information to identify DVGs and TVGs. STAVAG effectively identifies biologically meaningful DVGs for uncovering prognostically relevant gene modules that align with the malignant-immune-stroma structure in the tumor microenvironment, identifying marker genes in three-dimensional brain data, and detecting 12 polarization genes regulating planarian regeneration. Furthermore, by tracking the progression of myocardial infarction and tissue development, STAVAG reveals critical TVGs that can serve as diagnostic biomarkers for the ultra-acute phase of myocardial infarction. STAVAG also identifies both global and local gene modules involved in mouse placentation, as well as tissue-specific and interactive TVG modules that dynamically evolve during mouse embryonic development.

## Introduction

Spatial transcriptomics (ST) combines transcriptomic profiling with spatial information within a tissue. This technology enables us to examine gene expression and cell composition at specific locations, facilitating understanding of the microenvironment of malignant tissues^1^ and tracking spatial dynamics of gene expression during the formation of organs and the progression of diseases^2–4^. Various ST technologies are rapidly evolving, focusing on improving the resolution, sensitivity, and scalability^5^. Meanwhile, many computational methods have been developed to decipher spatial transcriptomics data, such as STAGATE, IRIS for the detection of spatial domains^6, 7^, STALocator, CellTrek for the integration of single-cell RNA-seq and ST data^8, 9^, STAGE and ST-Net for the generation of high-resolution ST profiles^10, 11^, STAMapper for the annotation of single-cell ST data^12^, and COMMOT, CellChat for the inference of spatial cell-cell interactions^13, 14^.

A key challenge in spatial transcriptomics applications is identifying genes specifically activated with spatial patterns, commonly referred to as spatial expression genes (SE genes) or spatially variable genes (SVGs). Identifying SVGs helps understand the spatial heterogeneity in tissues and the function of complex biological systems^15, 16^. However, beyond the foundational concept of spatial heterogeneity captured by SVGs, recent advances in spatial-temporal transcriptomics now enable us to probe deep biological questions through directional and temporal dimensions. For instance, given the direction of cancer invasion, we aim to explore variable genes along the tumor invasion axis^17^. Planarians have powerful regenerative capabilities and can reconstruct the correct anterior-posterior (AP) axis polarity during regeneration^18^. Therefore, identifying variable genes along the AP axis is crucial for uncovering key genes involved in AP axis polarization. Tracking variable genes during embryonic development can reveal how spatial patterns evolve to establish tissue structures^4^. Similarly, monitoring variable genes during disease progression could help identify key regulators of pathological changes^3^. To address these aspects, we introduced the concepts of directionally variable genes (DVGs) and temporally variable genes (TVGs). Here, DVGs refer to variant genes along a specific biological direction, while TVGs refer to genes that undergo dynamic changes over time.

Currently, existing methods for identifying gene expression patterns focus on SVGs. For example, SpatialDE applies a likelihood ratio test to assess whether the spatial covariance matrix effectively explains the gene expression in a Gaussian process regression^19^, nnSVG uses nearest neighbor Gaussian process models to estimate a gene-specific length scale parameter^20^, SPARK-X adopts a non-parametric covariance testing framework incorporated with various spatial kernels^21^, HEARTSVG excludes non-SVGs by testing the serial autocorrelations in the marginal expressions across the global space^16^. These methods have indeed been widely applied to conventional two-dimensional spatial transcriptomics. However, they are not capable of identifying DVGs or TVGs. This limitation poses a significant challenge in understanding the dynamic gene expression patterns and their role in various biological processes. Most of these methods are not directly applicable to three-dimensional (3D) data, highlighting a significant limitation given the real 3D biological systems. This limitation has become increasingly apparent with the rapid advancements in studying the 3D structures of the brain, cancer, and planarian^2, 22, 23^. Moreover, these methods have been predominantly tested on brain tissue samples, as there is insufficient evidence to support their applicability to other complex biological systems.

To this end, we propose a <u>s</u>patial or temporal-<u>a</u>ided framework for identifying directionally and temporally <u>va</u>riable <u>g</u>enes (STAVAG). STAVAG is a method based on the gradient boosting regression technique, designed to identify genes having both spatial and temporal patterns in a unified framework. STAVAG utilizes gene expression data to fit physical or temporal coordinates and evaluates the contributions of genes along any given dimension (either physical or temporal direction) based on their information gain. We demonstrate the benefits of STAVAG via applications to six different spatial transcriptomic data, which were derived from human cutaneous squamous cell carcinoma (cSCC)^17^, mouse cortex^24^, mouse hypothalamic preoptic region^25^, planarian^2^, heart affected by myocardial infarction (MI)^3^, mouse placentation development^26^, and mouse embryo development^4^. These data were obtained from diverse spatial transcriptomics technologies, including 10X Visium, STARmap, MERFISH, and Stereo-seq, demonstrating the broad applicability and robustness of STAVAG across different platforms.

## Results

### Overview of STAVAG

We illustrate the module schematic of STAVAG (**Fig. 1**, **Methods**). For a given spatial transcriptomics dataset, each spot or cell has a spatial coordinate (as well as a temporal coordinate for some scenarios). Here, we use **Y** to denote the spatial coordinates along an arbitrary direction in a 2D or 3D space, and, in a spatiotemporal context, can also indicate temporal information. We hypothesize that if a gene exhibits a directional or temporal pattern, its expression **X** should demonstrate a strong relationship to the direction or temporal order. STAVAG employs a gradient-boosting tree for regression to model the nonlinear relationship between the gene expression **X** and its **Y** coordinate (or temporal order) of each spot (or cell). For each direction, the importance of each gene is assessed based on the information gain it contributes to the regression task. STAVAG calculates the p-values by comparing the observed gene contribution score against the null distribution obtained through a single permutation (**Methods**). STAVAG determines DVGs and TVGs based on the information gained along a specified spatial direction or temporal order. STAVAG can also identify SVGs and demonstrates superior performance compared to existing methods (**Supplementary Note 1**).

**Figure 1.**
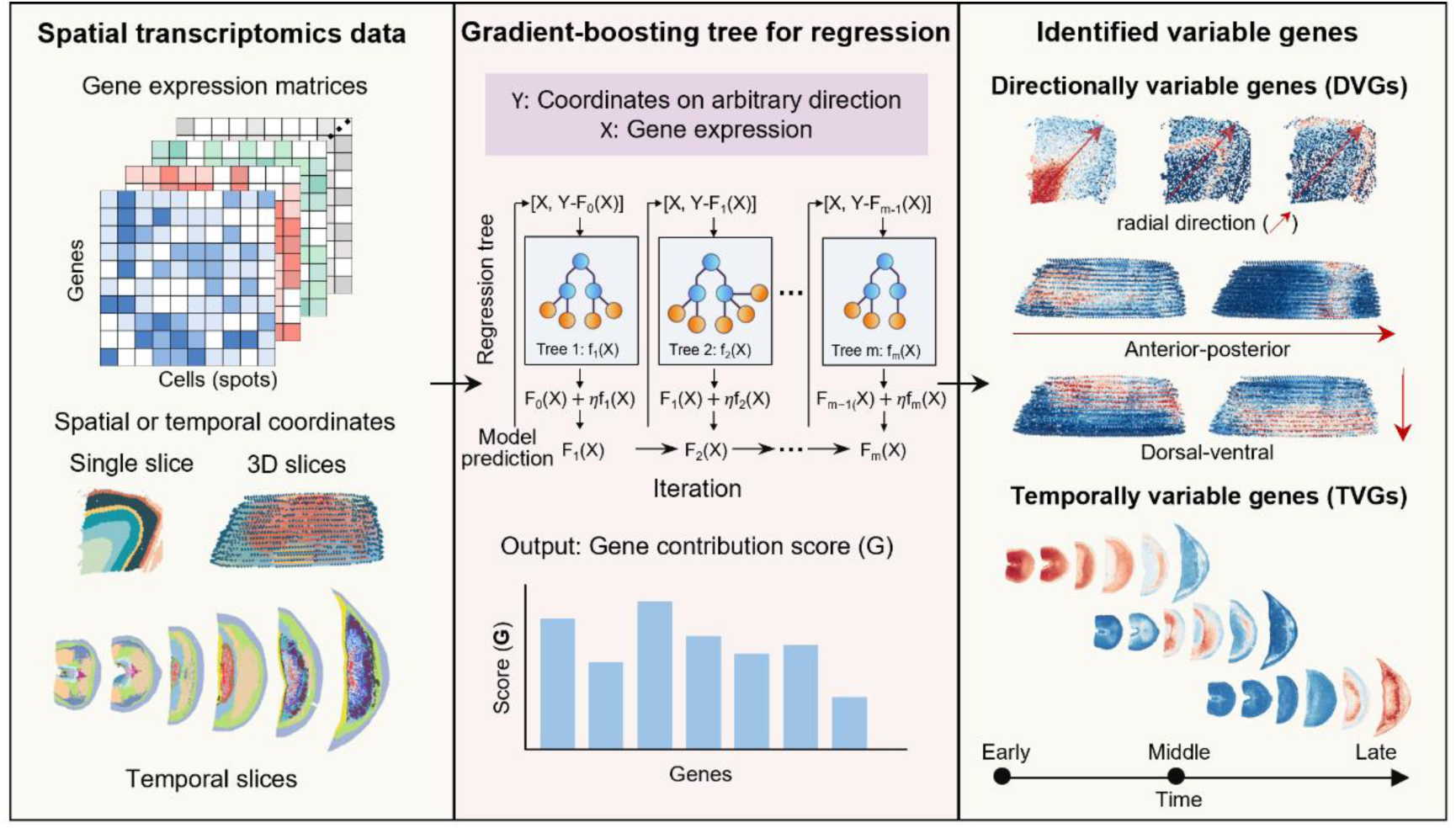
Illustration of STAVAG. STAVAG can handle spatial transcriptomics data for single, 3D, or multiple slices coupled with temporal information. STAVAG takes spatial transcriptomics data as input and then fits the spatial or temporal direction of spatial data with gene expression using a gradient-boosting tree for regression. STAVAG calculates the gene contribution score along any given direction or temporal order for each gene and identifies directionally variable genes (DVGs), and temporally variable genes (TVGs) for different scenarios.

### STAVAG identified DVGs that revealed the hierarchical microenvironment in cSCC data

The hierarchical tumor microenvironment is a hallmark of tumors^27^. Here, we applied STAVAG to a 10x ST dataset collected from cSCC tissue^17^. The cSCC data exhibit a top-to-bottom hierarchical structure along the y-axis, with the tumor region positioned at the top, followed by the adjacent normal region (**Fig. 2a, Supplementary Fig. 3a**). We also applied the cell-type deconvolution by RCTD as a reference (**Fig. 2a, Methods**)^17, 28^. Specifically, the tumor-specific keratinocytes (TSKs) and tumor cells were localized at the leading edge of the cancer region. In contrast, the immune cells, i.e., B cells, T cells, and Macrophages (Macro), were localized in the middle of the cancer region. The endothelial cells (EC) and fibroblasts were localized at the bottom of the cancer region (**Supplementary Fig. 3b**). Moreover, tumor_diff and tumor_basal display spatial distributions that are predominantly clustered toward the right end.

**Figure 2.**
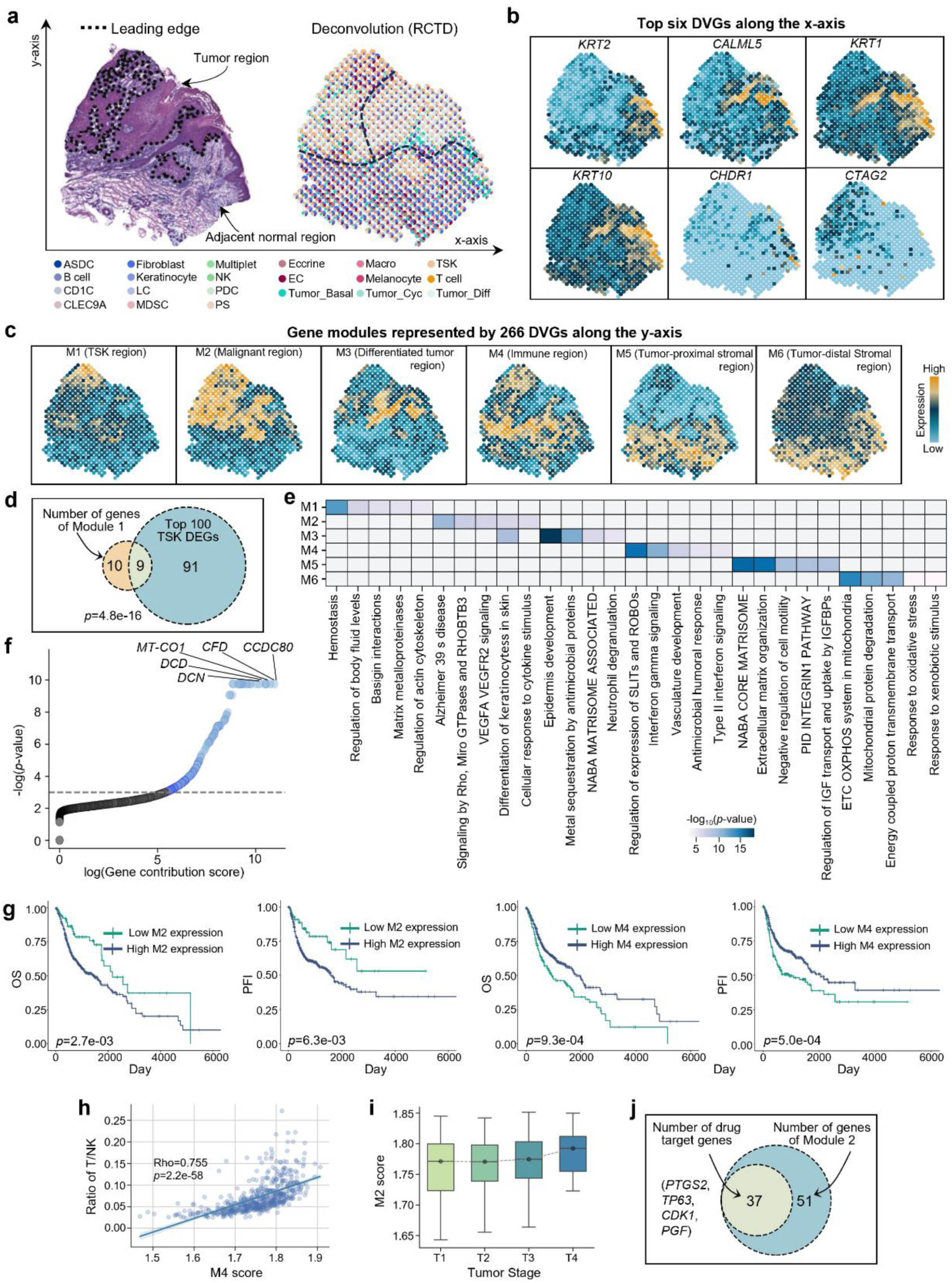
Analyzing the human cSCC ST data. **a.** Left, H&E staining with the leading edge of the tumor (dotted lines). Right, the deconvolution results of cSCC data using RCTD. Tumor_cyc (tumor cycling cell), tumor_diff (tumor differentiated cell), tumor_basal (tumor basal cell), EC (endothelial cell), PS (pilosebaceous)**. b.** The expression of six representative DVGs along the x-axis. **c.** The expression of gene modules aggregated from 266 DVGs along the y-axis. **d.** Venn plot showing the intersection of M1 genes and the top 100 TSK genes. The p-value was calculated using a two-sided hypergeometric test. **e**. Top five terms enriched for modules M1 to M6. Some pathway names were abbreviated; the full names can be found in **Table S1**. **f.** Gene contribution score versus significance of STAVAG along the y-axis for all genes in the cSCC data. **g.** Kaplan-Meier analysis showing the overall survival (OS) and progression-free interval (PFI) rates of patients, characterized by either low or high expression of M2 and M4 in the TCGA HNSC dataset (n = 566). The p-value was calculated using a two-sided log-rank test. **h.** Scatterplot showing the correlation between the fraction of T/NK cells and M4 score in the TCGA HNSC dataset. **i.** Boxplot showing the M2 score across tumor stages in the TCGA HNSC dataset. **j.** Venn plot showing the intersection of M2 genes and the drug target genes catalogued in DGIdb (https://dgidb.org/). The P value was calculated using a hypergeometric test.

To explore the spatial asymmetry of tumor cell distribution along the x-axis, we applied STAVAG in that dimension. STAVAG identified several DVGs along the x-axis, e.g., *CTAG2* on the left side of the cancer region, which is associated with tumor_cyc cells. In contrast, genes like *KRT1*, *KRT2*, *KRT10*, and *CALML5* are highly expressed in tumor_diff cells (**Fig. 2b**). Notably, both KRT genes and *CALML5* have been validated to be associated with the terminal differentiation of keratinocytes^29^. More importantly, the y-axis aligns with the tumor invasion gradient. We then applied STAVAG to this dimension and identified 266 DVGs, which were classified into six distinct modules, revealing a hierarchical spatial organization (**Fig. 2c, f, Supplementary Fig. 3c**). Notably, M1 (Module1) exhibited predominant expression in TSKs. Enrichment analysis revealed that their functions are associated with key TSK-related pathways, including matrix metalloproteinases, regulation of actin cytoskeleton, and cancer pathways^30^ (**Fig. 2e, Table S1**). Compared to the top 100 TSK differentially expressed genes identified in the single-cell analysis of cSCC data^17^, 9 out of 19 genes in M1 overlapped (p-value = 4.8e-16, **Fig. 2d**). These nine genes, characterized by both differential expression and distinct spatial patterns, are key TSK genes. Specifically, *IL24* functions during an inflammatory response in the skin by inhibiting the proliferation and migration of keratinocytes^31^; *SERPINE1* stimulates keratinocyte adhesion^32^; and *CAV1* regulates TSK proliferation^33^. Modules 2 and 3 are predominantly located in the upper-middle region of the section, primarily corresponding to the areas occupied by tumor cells. Both modules are enriched in the function of differentiation of keratinocytes in the interfollicular epidermis in mammalian skin, with a higher enrichment score observed in M3 (Differentiated tumor region), which is a hallmark of cSCC (**Fig. 2e**). In the central region of the slice, 66 genes from M4 are predominantly expressed. This region primarily corresponds to the location of immune cells (**Supplementary Fig. 3b**). It is associated with immune-related functions, including interferon gamma signaling, antimicrobial humoral response, and type II interferon signaling^34^. More importantly, high M2 expression is linked to poor survival, and high M4 expression is associated with better outcomes in the external TCGA HNSC dataset, which is another subtype of squamous cell carcinoma (**Fig. 2g**)^35^. Also, M4 is tightly linked to immune infiltration, showing a strong positive correlation with T/NK cell fractions (Spearman’s ρ = 0.755, p = 2.2 × 10⁻⁵⁸), and its high-expression stratum is consistently associated with favorable prognosis (**Fig. 2h**). In contrast, M2 increases monotonically with tumor stage, capturing progression biology; among its 88 constituent genes, 37 already have drugs catalogued in DGIdb, including validated cSCC targets such as *PTGS2*, *TP63*, and *CDK1*, while *PGF* has been implicated across multiple cancers (**Fig. 2i, j**).

We identified two distinct stromal modules, M5 and M6, mainly located beneath the tissue slices and enriched in stromal cells such as endothelial cells (EC) and fibroblasts (**Supplementary Fig. 3b**). Notably, M5 in tumor-proximal stromal regions, is characterized by strong enrichment of extracellular matrix, e.g., extracellular matrix organization, NABA core matrisome, PID Integrin1 Pathway. Additionally, these genes are enriched in the regulation of insulin-like growth factor (IGF) transport, a signaling axis critical for tumor-stromal interactions by promoting fibroblast activation and angiogenesis^36^. In contrast, module M6, present in tumor-distal stroma, is enriched for integrin-mediated signaling pathways, reflecting a more homeostatic microenvironment with intact cell-ECM interactions and limited structural remodeling.

In short, STAVAG deciphered the tumor-immune-stroma microenvironment in cSCC, and the gene modules it identified accurately represent the functional characteristics of each region. Moreover, STAVAG identified specific modules associated with poor prognosis, offering potential biomarkers for risk stratification and therapeutic targeting. Notably, the majority of these DVGs are also SVGs, as they exhibit strong spatial expression patterns. However, SVGs represent an overbroad concept that does not inherently capture directionality or timeliness, making it challenging to extract finer biological insights.

### STAVAG identified DVGs along diverse spatial directions within 3D data

3D spatial transcriptomics captures the true spatial architecture and has recently gained significant attention^22, 23^. We here utilized a 3D section of a STARmap mouse cortex data with nine manually annotated cell types^24^, which consists of 89 consecutive slices (**Fig. 3a**).

**Figure 3.**
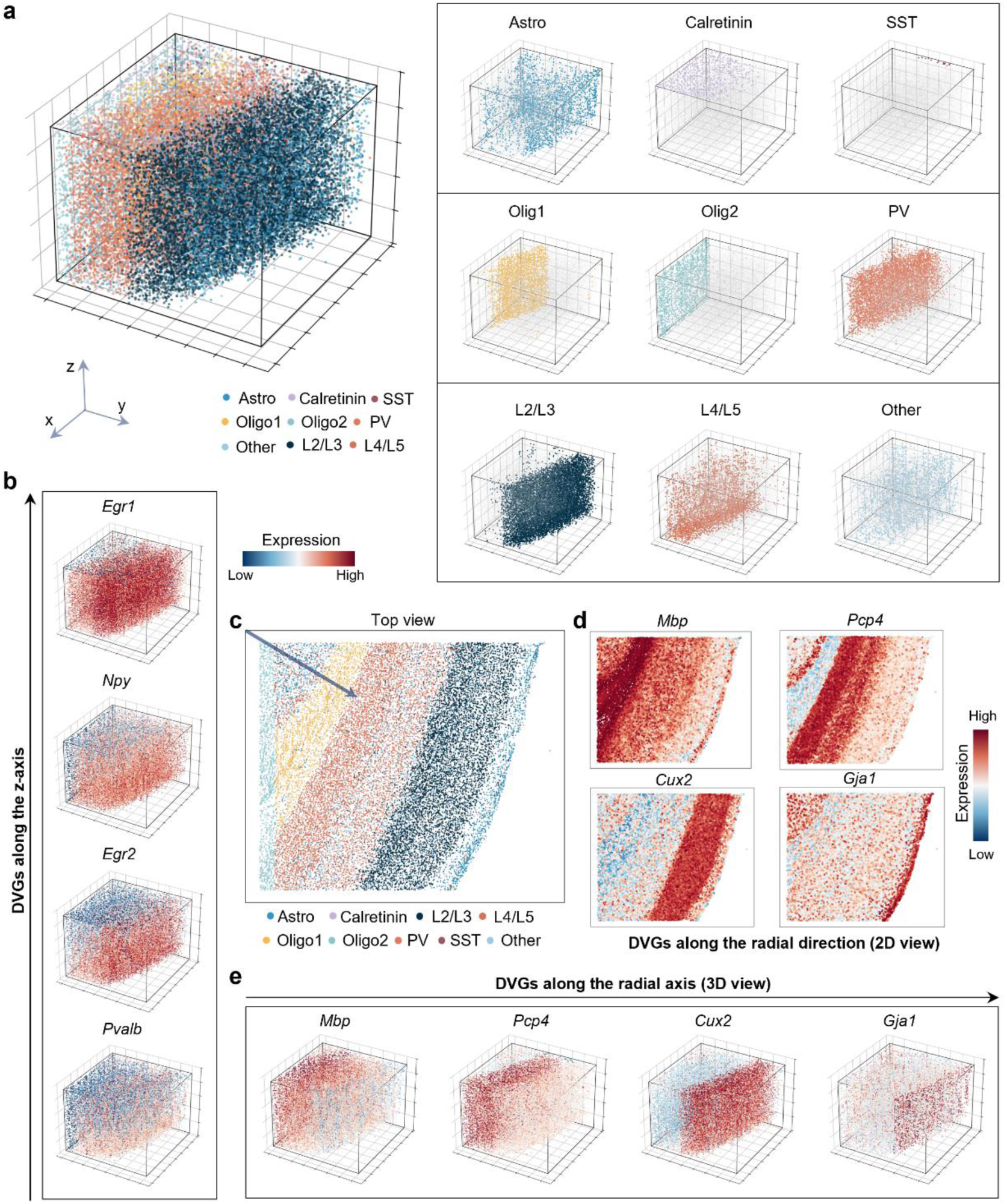
Analyzing the STARmap mouse cortex data. **a.** Overview of the STARmap mouse cortex dataset with its annotated cell types. **b.** The expression of DVGs along the z-axis with different expression patterns. **c.** The 2D view of mouse cortex data, related to (**a**), shows arrows pointing diagonally downward at a 30° angle, indicating the direction in which STAVAG identifies DVGs. **d-e.** The 2D (**d**) and 3D (**e**) views illustrate the expression patterns of DVGs along the specified direction.

In 3D spatial transcriptomics, the z-axis adds dimension to 2D, thereby providing more comprehensive information. By applying STAVAG to this axis, we identified several DVGs, e.g., *Pvalb*, *Egr2*, *Npy*, and *Egr1*, which exhibit a progressively increasing expression trend along the z-axis. Specifically, *Pvalb* is primarily expressed in the lower region, and *Egr1* is expressed throughout all regions except for the top (**Fig. 3b**). Besides, we applied STASVG to the z-axis of MERFISH hypothalamic preoptic region data^25^ (**Supplementary Fig. 4a**). We identified numerous cell-type markers whose expression patterns were influenced by significant variations in cell numbers along the z-axis. Notable examples include *CD24a*, *Mbp*, and *Sln*, which exhibited high expression changes corresponding to these variations. Additionally, we observed that *Crh* was exclusively expressed in the bottommost slice, which may be related to the existence of GABAergic interneurons^37^ (**Supplementary Fig. 4b**).

The cortex exhibits a hierarchical organization from the inner to the outer layers, with distinct cellular compositions and functional specializations at each level^38^ (**Fig. 3c**). Next, we applied STAVAG to the radial direction of the data, aiming to identify DVGs associated with the hierarchical organization. This approach revealed numerous potential markers consistent with the cortical hierarchical organization, such as *MBP*, which is primarily expressed in Oligo1; *PCP4*, predominantly found in PV and L4/L5; *Cux2*, mainly expressed in L2/L5; and *Gja1*, which is primarily localized in Astro (**Fig. 3d, e**).

In summary, STAVAG provides a flexible framework for identifying DVGs along biologically meaningful directions and performs effectively in 3D spatial transcriptomics. Notably, *Pcp4* exhibits spatial variability (**Fig. 3d, e**) and is identified as a DVG along the radial direction but not along the z-axis (**Supplementary Fig. 5a, b**). This distinction highlights the critical role of directional considerations in identifying spatially variable genes, emphasizing that variability may be direction-dependent rather than uniform across spatial dimensions.

### STAVAG identified DVGs that uncovered the AP axis polarization on 3D planarian data

We next analyzed a 3D section of a planarian from our previous study, which represents an entire planarian body and consists of twelve 2D 10X Visium spatial transcriptomic slices^2^. It includes ten manually annotated spatial domains, revealing the tissue and cell distribution map of the planarian (**Fig. 4a, Supplementary Fig. 6a**).

**Figure 4.**
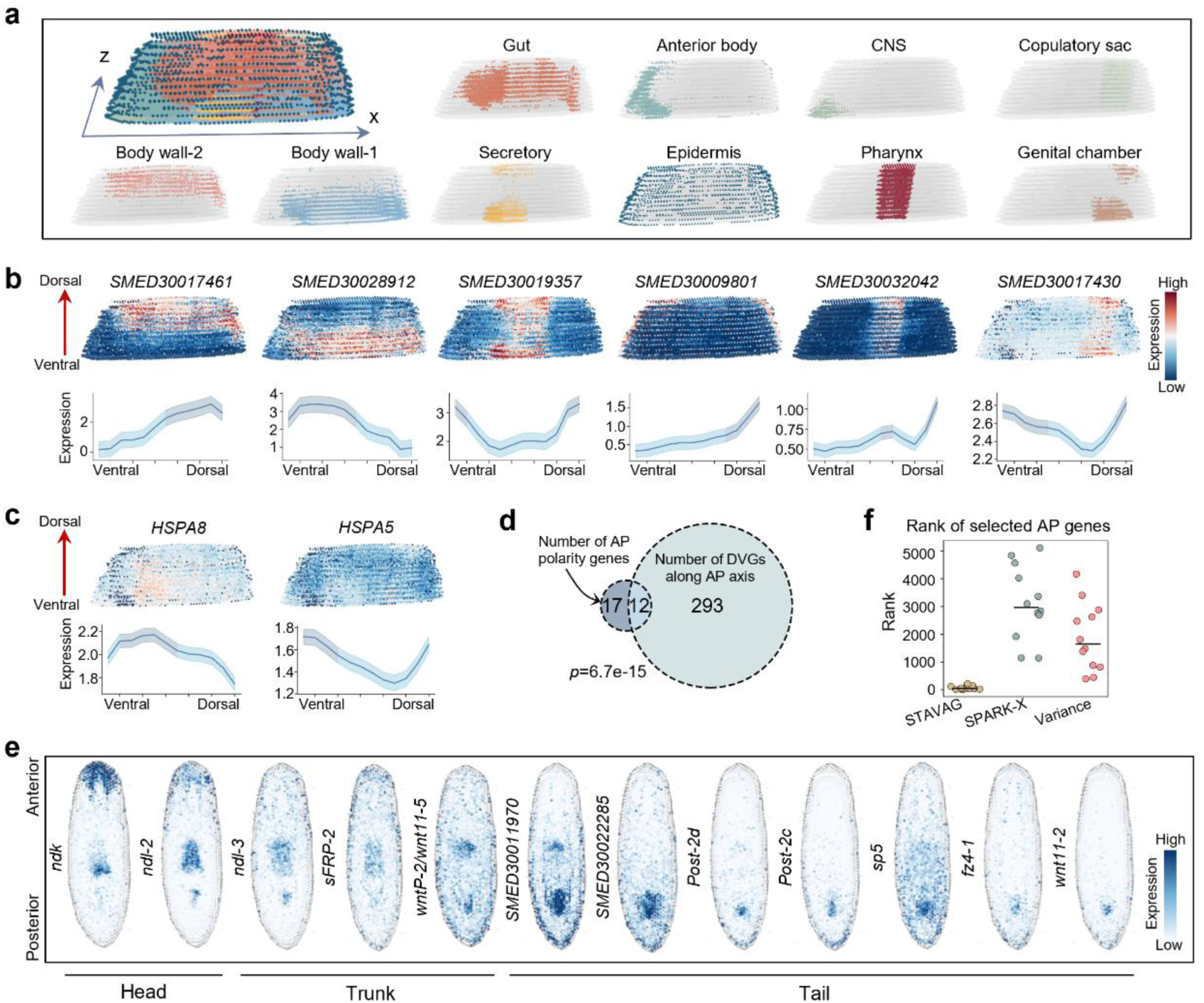
Analyzing the 3D planarian ST data. **a.** Overview of the planarian dataset with its annotated spatial domains. **b** and **c.** The expression of DVGs along the DV (z) axis with different expression patterns. **d.** Venn plot showing the intersection of AP polarity genes and DVGs along the head-tail axis. The *p*-value was calculated using a two-sided hypergeometric test. **e.** Top-view visualization showing the expression of 12 AP genes. The location marked by the horizontal line denotes the regeneration site under their regulation. **f**. Comparison of ranks of 12 AP genes identified by STAVAG, SPARK-X, and Variance, respectively, with the horizontal line indicating the median value.

The distribution of tissue types along the dorsal-ventral (DV) axis of planarians exhibited a significant hierarchical organization^2^. Additionally, several key wound-induced genes associated with regeneration were found to be highly expressed in specific tissue^18^. We then searched for DVGs along the DV axis and identified numerous genes that are consistent with the distribution of spatial domains and could serve as potential marker genes (**Fig. 4b, Supplementary Fig. 6b, c**). Notably, these genes often exhibit distinct patterns of increase or decrease, or even an initial decrease followed by an increase, along the DV axis, highlighting the potential of STAVAG to identify DVGs with diverse spatial patterns. In addition, we also identified two heat shock proteins, *HSPA8* and *HSPA5*, both of which are associated with wound-induced responses^18^ (**Fig. 4c**).

Regeneration is a key characteristic of planarians, allowing them to regrow lost tissues or organs after injury. In particular, the AP axis polarization genes that control head, trunk, and tail regeneration have garnered significant attention, as RNAi knockdown of these genes can lead to AP axis polarization defects such as posterior-facing heads and the formation of two pharynges^39^. Along the AP axis, we discovered 305 DVGs, excitingly including 12 of the 29 AP axis polarization genes (p-value = 6.7e-15, **Fig. 4d, e**). The remaining 17 genes exhibited very low expression levels (median value = 0.023) and were therefore not detected by STAVAG (**Supplementary Fig. 7a**). Notably, these 12 AP axis polarization genes ranked highly in the STAVAG analysis and exhibited constitutive regional expression patterns along AP axis (**Supplementary Fig. 8**). As a comparison, SPARK-X, which identifies SVGs without considering directions, could not effectively identify the AP axis polarization genes within the top SVGs (median rank = 2968) (**Fig. 4f, Supplementary Fig. 8**). We also tried a simple method to address this task. Following the approach of the original study, we divided the AP axis into ten equal segments. We then attempted to calculate the variance of each gene along these segments to identify AP axis polarization genes (**Methods**). Although this approach showed a slight improvement over SPARK-X (median rank = 1647.5), it still did not effectively capture the AP axis polarization genes. Moreover, STAVAG clusters all DVGs into distinct gene modules, revealing their differential spatial distribution along the AP axis (**Supplementary Fig. 9a, Table S2**). Notably, we identified three gene modules enriched in the head, trunk, and tail, respectively, which collectively encompass nearly all AP polarity genes (11 out of 12) (**Supplementary Fig. 9b**). Therefore, through module analysis, STAVAG demonstrates an enhanced capability to identify potential AP polarity genes specific to designated regions, providing valuable guidance for downstream biological experiments.

In conclusion, STAVAG not only performs exceptionally well in 3D spatial transcriptomics but also facilitates the identification of DVGs with significant biological relevance, offering reliable targets for subsequent biological experiments. We also showed that STAVAG outperforms other competing methods in identifying SVGs in 3D datasets, highlighting its effectiveness and versatility in spatial transcriptomic analysis. (**Supplementary Notes 4**).

### STAVAG identified key signals involved in the progression of myocardial infarction

Acute myocardial infarction (AMI) is the primary cause leading to human mortality and morbidity. Rapid diagnosis in the ultra-acute phase is of high clinical value; millions of patients present with chest pain each year, and minutes lost before diagnosis translate directly into worse outcomes^40, 41^. We next applied STAVAG to 10x ST datasets from post-MI samples collected at 1, 4, 72, and 168 hours^3^. All spots were annotated as remote zones (RZ), border zone 1 (BZ1), BZ2, and infarct zones (IZ) regions, following a hierarchical structure (**Fig. 5a**). RZ represents relatively normal myocardium and is separated from the cell death (IZ) by the BZ zone. Our goal is to discover potential biomarkers that can facilitate early and more accurate diagnosis of AMI.

**Figure 5.**
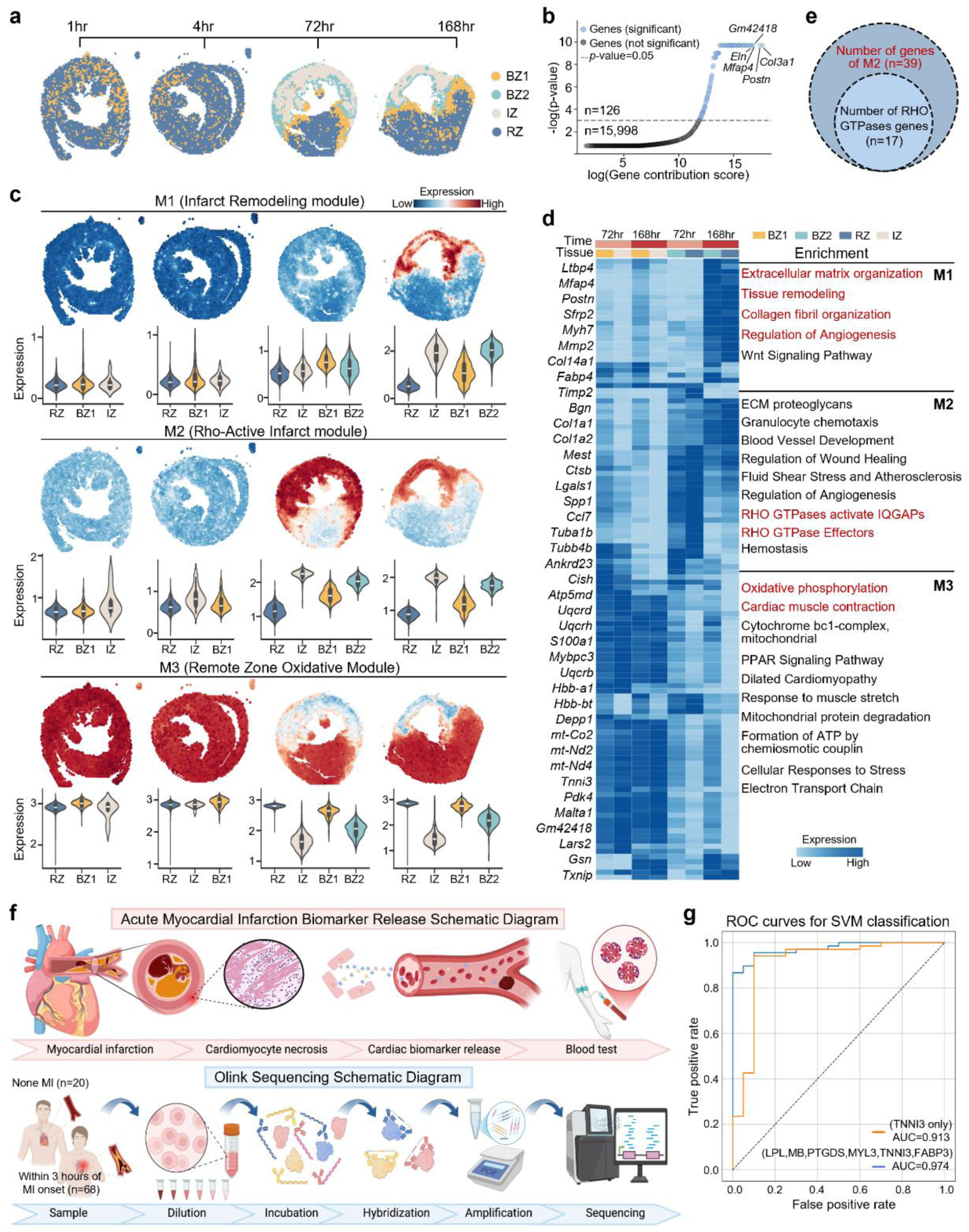
Analyzing the mouse myocardial infarction ST data. **a.** Overview of the myocardial infarction dataset with its annotated spatial domains **b**. Gene contribution score versus the significance of STAVAG for all genes in the mouse myocardial infarction data. **c**. Dotplot showing the expression of three TVG modules on spatial data. Violin plot showing the expression of three TVG modules on each domain. **d.** Heatmap showing the expression of TVGs and the corresponding enrichment terms. **e.** Venn plot showing the number of RHO GTPases genes in M2. **f.** Illustration of acute myocardial infarction and the Olink sequencing process. **g.** Receiver operating characteristic (ROC) curves, derived from SVM classification, comparing TNNI3 alone with the six predictive proteins identified in M3.

To understand the temporal dynamics underlying MI progression and repair, we applied STAVAG to the MI data along the temporal axis and identified 126 TVGs. Among the top five significantly expressed genes, *Col3a1*, *Postn*, *Mfap4*, and *Eln* are associated with cardiac and vascular stability, as well as extracellular matrix repair processes^42–44^, thereby validating the effectiveness of STAVAG (**Fig. 5b**). STAVAG clustered all TVGs into five gene modules, each exhibiting enrichment at different temporal stages (**Supplementary Fig. 10a**). M1 starts with minimal levels at early time points (1 and 4 hours), gradually rising over time and reaching its peak at 168 hours. Its expression is predominantly elevated in the IZ and BZ2 regions, particularly at later time points, while the RZ maintains consistently lower expression levels throughout (**Fig. 5c**). Genes within M1 are enriched for processes associated with extracellular matrix organization, tissue remodeling, collagen fibril organization, and regulation of angiogenesis, suggesting their involvement in structural repair and vascular development (**Fig. 5e**). These findings indicate that M1 plays a critical role in facilitating tissue remodeling and angiogenic responses, especially within the IZ and BZ2 regions, which are likely key sites of active repair following injury. M2 also shows a time-dependent increase, with low expression at 1 and 4 hours and a significant rise in IZ and BZ2 at 72 and 168 hours, unlike M1, which lacks high IZ expression at 72 hours (**Fig. 5c**). Among the 39 genes in M2, we identified 17 genes associated with RHO GTPases, which were significantly enriched for RHO GTPase-related functions (**Fig. 5e, d**). This finding is consistent with the original conclusion that RHO GTPase signaling was considered a critical function in late-stage developing cardiomyocytes^3^. In contrast to Modules 1 and 2, M3 displays a stable expression pattern in RZ and BZ1 regions. The genes in M3 are enriched for pathways related to oxidative phosphorylation, cardiac muscle contraction, and electron transport chain, highlighting its role in maintaining energy production and metabolic homeostasis (**Fig. 5e**). This suggests that normal myocardium supports the high energetic demands of repair and regeneration processes by ensuring cellular energy supply and stress adaptation, which are essential for sustaining tissue function and recovery. Also, we identified TVGs in RZ regions (**Supplementary Note 5**).

Importantly, we would like to test, in human patients, whether proteins encoded by M3 genes can predict MI in the ultra-acute window. We externally validated M3 in an independent cohort of patients presenting with chest pain within 3 hours of symptom onset (n = 88), with all data prospectively collected by our research team (**Methods**). Subsequent coronary angiography and clinical adjudication confirmed AMI in 68 patients, whereas 20 were confirmed to have no severe coronary artery stenosis. We profiled plasma using an Olink proximity extension assay to quantify 729 proteins; six of these proteins (LPL, MB, PTGDS, MYL3, TNNI3, FABP3) overlapped with the M3 gene set. We then evaluated this six-protein M3 signature and found that it accurately predicts early AMI. Using a support vector machine with leave-one-out cross-validation, the M3 composite signature (which includes the canonical marker TNNI3) achieved an AUC of 0.974, substantially outperforming TNNI3 alone (AUC = 0.913). Moreover, rather than merely picking the most discriminative proteins from the Olink panel, we leveraged STAVAG-derived TVGs that we independently confirmed are highly expressed in the heart during the ultra-acute phase of MI, thereby conferring greater biological (and disease) specificity. Collectively, these findings highlight STAVAG’s capacity to move from computational module discovery to clinically actionable diagnostic biomarkers.

### STAVAG identified global and local gene modules involved in the development of mouse placentation

We next applied STAVAG to a high-resolution ST dataset consisting of 102,286 cells on mouse placentation spanning from embryonic day E7.5 to E14.5 profiled by Stereo-seq^26^. The annotated regions and subregions from the original analysis depict the spatiotemporal landscape of the placentation (**Fig. 6a, Supplementary Fig. 11a**).

**Figure 6.**
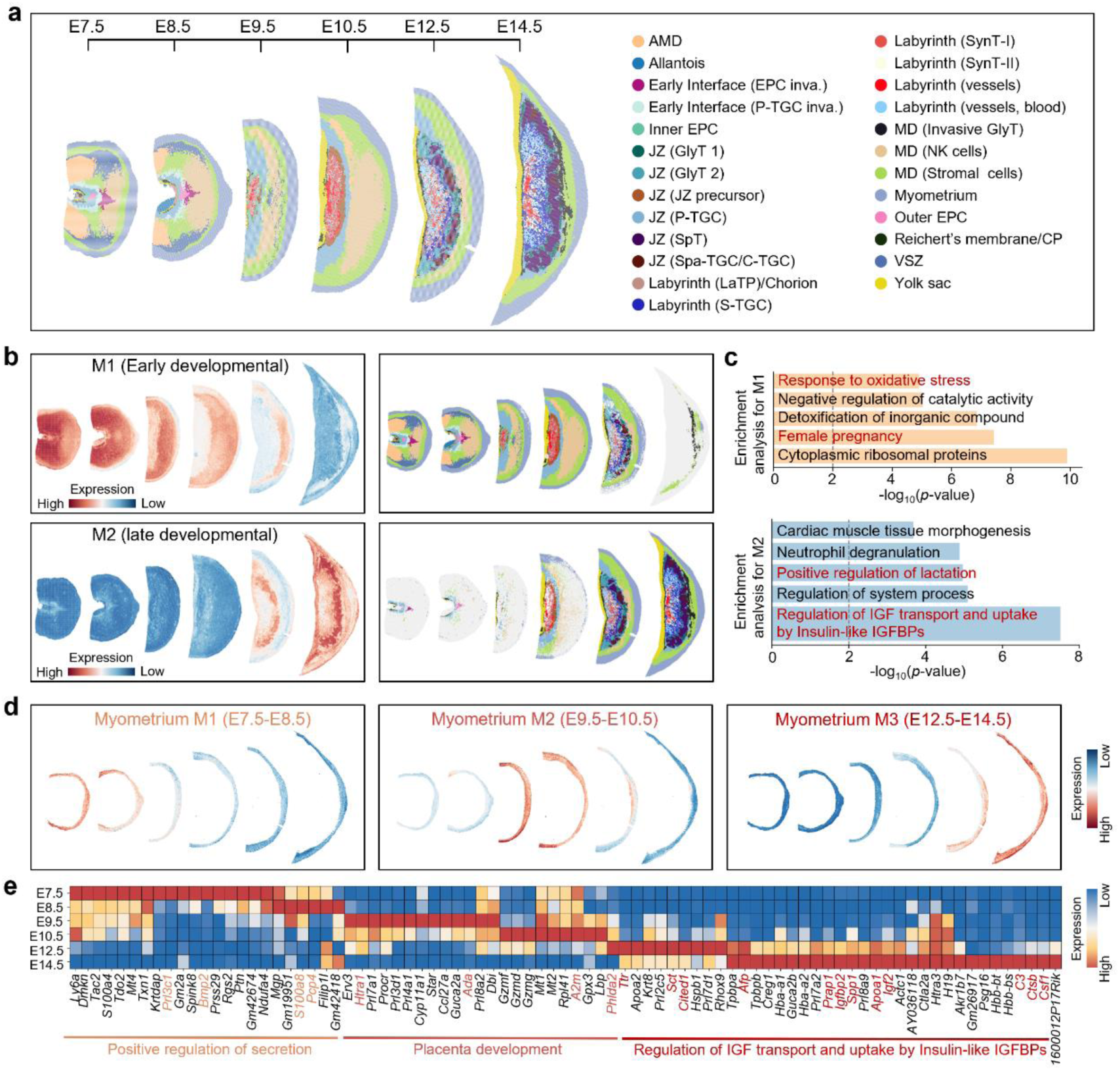
Analyzing the mouse placentation Stereo-seq data. **a.** Overview of the mouse placentation dataset with its annotated spatial sub-domains. **b.** Dotplot showing the expression of two global TVG modules on spatial data (left panel) and their primary expression regions (right panel). **c.** Enrichment analysis of genes from M1 and M2. **d.** Dotplot showing the expression of three TVG modules identified from the myometrium. **e.** Heatmap showing the expression of TVGs identified in the myometrium as well as the corresponding enrichment terms. IGF (insulin-like growth factor), IGFBPs (insulin-like growth factor binding proteins).

STAVAG identified 148 TVGs associated with placental development, which were grouped into five distinct gene modules (**Fig. 6b, Supplementary Fig. 11b, c**). We identified two global gene modules, M1 and M2, which exhibit a complementary expression pattern, with M1 predominantly expressed in early developmental stages and M2 in later stages. M1 showed higher expression in regions that already exist at earlier stages, e.g., AMD, Early, and MD, with its expression gradually decreasing as development progresses. The enrichment analysis reveals several functions closely related to early placental development, including oxidative stress, which is crucial for managing the oxidative environment during early development and protecting cells from oxidative damage in the early placenta^46, 47^. The involvement in female pregnancy further underscores the module’s relevance to the foundational stages of placental formation and function (**Fig. 6c**). In contrast, M2 primarily localizes to regions such as the yolk sac and JZ (SPT), which emerge during the later stages of development (**Fig. 6b**). Enrichment analysis revealed its association with the positive regulation of lactation, a function closely linked to the later stages of placental development. Additionally, M2 is enriched in the regulation of IGF transport and uptake by Insulin-like IGFBPs, which is a critical regulator of placental growth and development^48^. Furthermore, we identified three additional regions exhibiting temporally dynamic changes within local spatial domains, underscoring the temporal relevance of the gene modules identified by STAVAG (**Supplementary Fig. 11c**).

Next, we focus on the TVGs in the myometrium region. The myometrium region is present throughout all stages of placental development, which makes us curious to investigate its temporal dynamics and explore its specific roles in this process (**Fig. 6d**). STAVAG identified 83 genes with dynamic expression patterns in the myometrium throughout placental development and grouped them into three distinct temporal modules (**Supplementary Fig. 12a**). Temporally, myometrium M1 peaks early and declines, M2 is active during mid-development, and M3 increases later, reflecting maturation. Functionally, myometrium M1 is associated with early processes such as the positive regulation of secretion, which represents one of the functional roles of the myometrium^49^. Myometrium M2 is linked to key events in placental development during mid-stage. In contrast, myometrium M3 is enriched in pathways related to IGH transport and metabolic regulation, which are crucial for regulating cell growth and differentiation in the later stages of placental development^50^ (**Fig. 6e, Supplementary Fig. 12b**). Moreover, previous studies have confirmed that *IGFBP2* is highly expressed in the myometrium^50^; however, our findings indicate that it is a gene whose expression increases only during the later stages in the development of mouse myometrium (**Fig. 6e)**.

Collectively, these findings highlight STAVAG’s capability to uncover global or local gene dynamics associated with the spatiotemporal progression of placental development, offering valuable insights into the key biological pathways at different stages of this process. We further down-sampled the data and conducted runtime tests, revealing a linear growth trend between the runtime and the number of cells (**Supplementary Fig. 11d**). This observation suggests that the method is scalable and well-suited for analysis on larger datasets.

### STAVAG identified tissue-specific and interactive functions during mouse embryonic development

Finally, we applied STAVAG to a Stereo-seq mouse embryos dataset, covering embryonic days 9.5 to 16.5 with one-day intervals^4^ (**Fig. 7a**). STAVAG identified 358 TVGs associated with embryonic development, which were grouped into 17 distinct gene modules (**Supplementary Fig. 13a, Table S3**). Among them, STAVAG identified tissue-specific ones (epidermis, GI tract, dorsal root ganglion, cartilage primordium, muscle, bone), whose expression progressively increased over time. They contain numerous marker genes and are potentially associated with the structural development or functional maturation of their respective tissue types (**Supplementary Fig. 13b**). We also observed the mutually exclusive expression patterns of M8 and M14. M8 is enriched in metabolically active organs such as the heart, liver, and bone. M14 genes are absent from these regions and may instead reflect regulatory programs in distinct lineages such as neural or germline tissues (**Supplementary Fig. 14a**).

**Figure 7.**
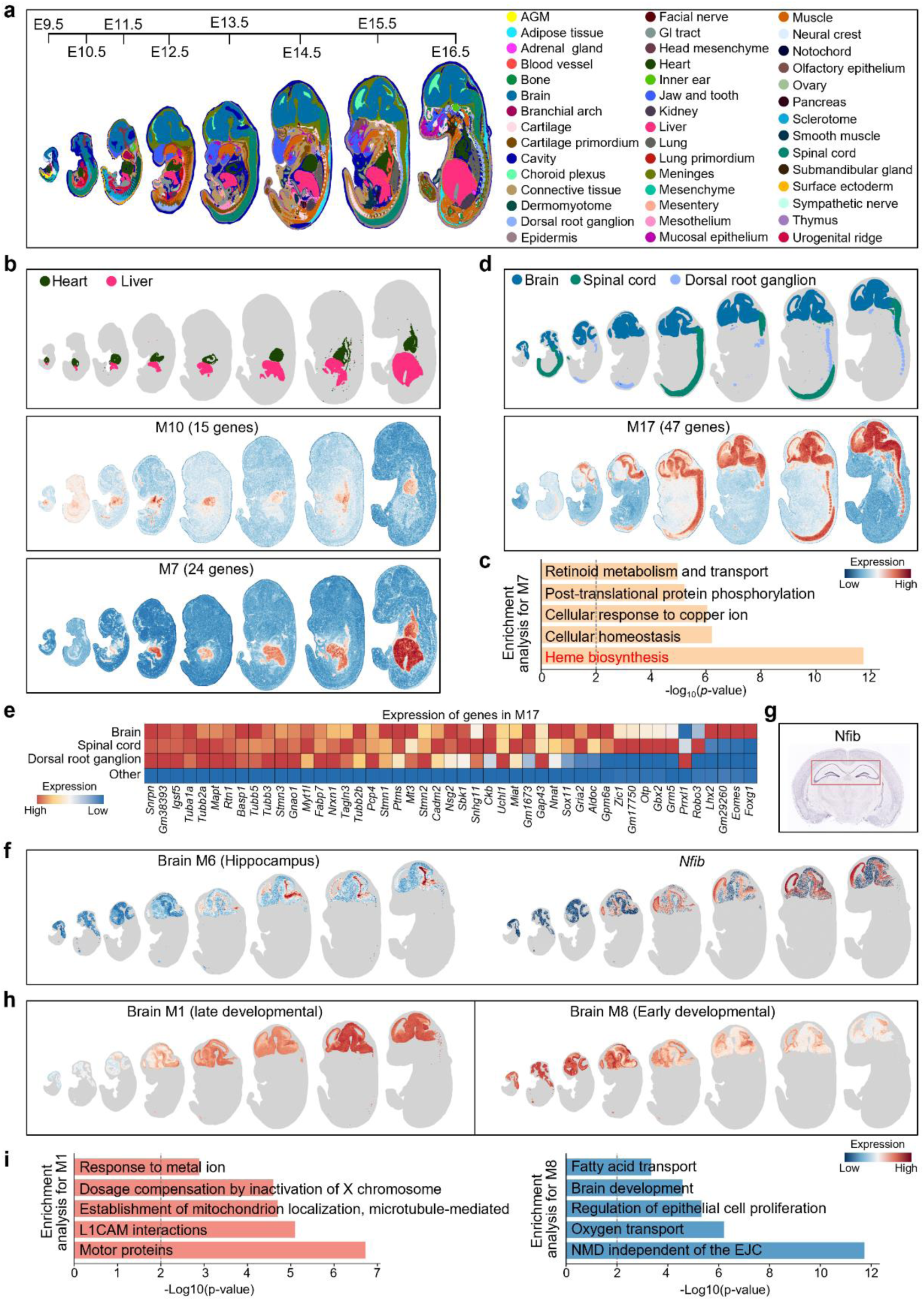
Analyzing the mouse embryonic development Stereo-seq data. **a.** Overview of the mouse embryonic dataset with its annotated spatial domains. **b.** Dotplot showing the location of the heart and liver and the expression of M7 and M10 on spatial data. **c.** Enrichment analysis of genes from M7. **d.** Dotplot showing the location of the brain, spinal cord, and dorsal root ganglion and the expression of M17 on spatial data. **e.** Heatmap showing the expression of genes in M17 grouped by different tissues. Other means all tissues except the brain, spinal cord, and dorsal root ganglion. **f**. The expression of brain M6 (left panel) and one representative gene (Right panel). **g**. ISH of Nfib from Allen Brain Atlas. **h.** The expression of one late and one early developmental brain module on spatial data. **i**. Top five terms enriched for modules M1 to M8.

Moreover, STAVAG identified a tissue-interactive module, M7, expressed in the heart and liver, which exhibited a delayed onset of expression in the heart, appearing only after embryonic day 16.5 (**Fig. 7b**). Before this stage, M7 was exclusively expressed in the liver, where it was significantly enriched for genes involved in heme biosynthesis (**Fig. 7c, Supplementary Fig. 14b**). This suggests that the liver may serve as a primary site of heme production during early development^51^, supplying essential heme-related factors to other developing tissues, including the heart. M17, identified by STAVAG, is expressed in the brain, spinal cord, and dorsal root ganglion at every developmental stage, with its expression gradually increasing over time (**Fig. 7d, e**). This progressive upregulation suggests an increasingly critical role of these organs in neural development and connectivity. This module is associated with key biological processes, e.g., neuron migration, neuron projection morphogenesis, and microtubule polymerization or depolymerization (**Supplementary Fig. 14c**), indicating its essential coordinating role in neurodevelopmental processes across these three tissues.

To understand the molecular mechanisms underlying brain development, we applied STAVAG to the brain and identified eight distinct TVG modules. We found that brain module M6 exhibits region-specific expression patterns and includes the gene *Nfib*, which, according to in situ hybridization (ISH) data from the Allen Brain Atlas, is predominantly enriched in the hippocampal region^52^ (**Fig. 7f, g**). Also, we found two modules representing distinct spatial patterns: M1 exhibits higher expression at later developmental stages, while M8 is more prominently expressed at earlier stages (**Fig. 7h**). Notably, their functional roles appear to be closely linked to their temporal expression patterns, enrichment analysis indicates that M8 may contribute to early brain formation, laying the foundation for neural structures. In contrast, M1, expressed at later stages, plays a role in refining neural circuits and functional specialization as development progresses (**Fig. 7i**). This highlights a dynamic, stage-specific regulation of neurodevelopmental processes orchestrated by these modules.

## Conclusion

Accurate modeling of spatial gene expression variation is a critical issue for understanding the intricate organization of tissues and organs. Numerous methods have been developed to address this^16, 19–21^. However, these methods fail to model gene expression changes along biologically meaningful dimensions such as a specific spatial direction or temporal progression.

In this work, we introduce the concepts of DVG and TVG and propose a tool, STAVAG, to identify them. DVG refers to variant genes along a given biological direction, such as the anterior-posterior axis in planarians, the direction of cancer invasion, or the boundaries of immune responses. TVG refers to genes that undergo dynamic changes, such as those associated with tissue development or disease progression. These genes are critical for advancing our understanding of the spatiotemporal dynamics of biological processes. STAVAG employs a gradient-boosting tree model to directly establish the relationship between gene expression and arbitrary directions or time directions, enabling the rapid identification of DVGs and TVGs. Unlike most algorithms limited to 2D spatial transcriptomic data^16, 19, 20^, STAVAG can be applied to 3D data or even 4D data that includes both spatial and temporal dimensions.

Without relying on external single-cell data, STAVAG successfully elucidates the hierarchical tumor-immune-stroma structure in cancer tissue slices and identifies two gene modules that are significantly associated with patient prognosis. Furthermore, in planarian data, STAVAG accurately identified 12 AP axis polarization genes. Moreover, it holds promise for guiding downstream experimental efforts aimed at discovering novel genes involved in AP polarity. In MI data, STAVAG precisely identified key gene modules associated with the temporal dynamics of disease progression. In mouse placenta development data, STAVAG detected gene modules enriched at distinct stages of development and linked these modules to specific developmental functions. During embryonic mouse development, STAVAG uncovered tissue-specific and interactive TVG modules, revealing how gene expression evolves to drive tissue development and coordinated interactions.

Due to its direct modeling approach and the algorithm’s simplicity, we believe that STAVAG provides an extensible framework. For example, it could integrate multiple methods to assess the importance of DVGs or TVGs. The gene interaction information can be incorporated as prior knowledge for STAVAG. Additionally, STAVAG can be easily extended to spatial data from other omics, such as spatial ATAC-seq data.

## Methods

### Data preprocessing

For all spatial transcriptomics datasets, we first filtered genes expressed in fewer than ten cells. Next, we performed library size normalization for each cell and applied a logarithmic transformation to the expression data with a pseudo-count.

### Architecture of STAVAG

STAVAG takes the preprocessed expression matrix 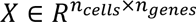 and the corresponding coordinate matrix 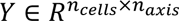 as input. Generally, the first and second dimensions of the coordinate refer to the x and y axes of ST data, while the third dimension represents the z-axis or the temporal axis. STAVAG applies the light gradient boosting machine^53^ to learn a spatiotemporal unified framework that maps the expression to its coordinates. Here, 𝑋_𝑖_ denotes the 𝑖 th row of 𝑋, corresponding to the expression profile of the 𝑖 th cell, and 𝑦_𝑖,𝑗_ denotes the element located at the 𝑖th row and 𝑗th column of 𝑌, referring to the 𝑗th coordinate of the 𝑖th cell. We repeatedly learnt 𝑚 boosting trees based on the error. Specifically, our goal is to use the ensemble of these boosting trees to minimize the mean squared error (MSE) on an axis 𝑗:

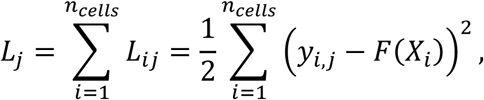

where 𝑗 ∈ {1, …, 𝑛_𝑎𝑥𝑖𝑠_}, and 𝐹(𝑋_𝑖_) denotes the output of the model. We minimize the loss function 𝐿_𝑗_ by iteratively adding weak learners such as trees to correct residuals, i.e.,

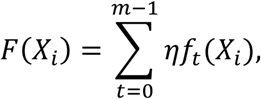

where 𝑓_𝑡_ denotes the boosting tree at iteration 𝑡 and 𝜂 is the learning rate. In the training procedure, 𝑓_𝑡_s are trained sequentially. 𝑓_0_(𝑋_𝑖_) is initialized as the average of 𝑦_𝑖,𝑗_ over all 𝑗, while at iteration 𝑡^∗^, the objective of a new boosting tree 𝑓_𝑡_∗(𝑋_𝑖_) is to minimize the residue of previous predictions:

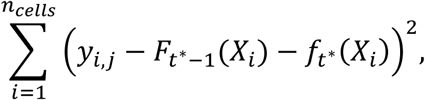

where 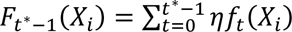 is the prediction from previous iterations.

In each iteration, decision trees are built by using a leaf-wise growth strategy, prioritizing splits that maximize the loss reduction. In detail, the values of each gene are discretized into histograms to accelerate split finding. We select the gene with the highest information gain to split the instance set and generate new leaves to the tree. The information gain of the gene 𝑘 is:

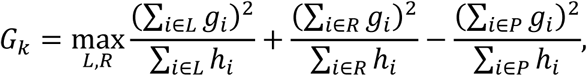

where the left child set 𝐿 and right child set 𝑅 are split from the parent set 𝑃 according to the expression of gene 𝑘 and 𝑃 = 𝐿 ∪ 𝑅. Here, 𝑔_𝑖_, ℎ_𝑖_ are the first and second order of gradient, i.e.,

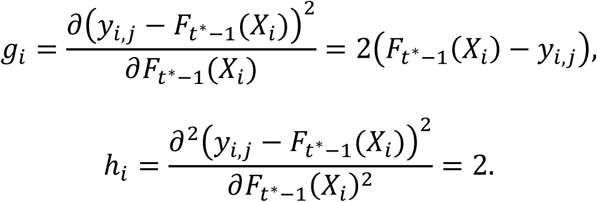

After training the model, we derive the STAVAG contribution score of the gene 𝑘 along the axis 𝑗 by aggregating the contributions from the learned boosting trees, which is defined as:

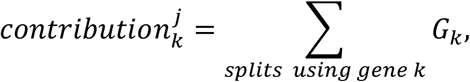

where 𝐺_𝑘_ is the information gain of gene 𝑘 in the split. To integrate the information from multiple axes, the aggregated STAVAG contribution score is calculated as follows:

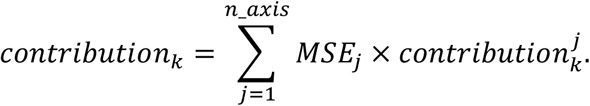

Here, 𝑀𝑆𝐸_𝑗_ denotes the MSE on the axis 𝑗. We utilize the STAVAG contribution score to quantify the contribution of the 𝑘 th gene’s expression concerning the spatial or temporal direction. We evaluated the importance of SVG by the aggregated STAVAG contribution score and the importance of DVG/TVG by the STAVAG contribution score along the given directional or temporal axes.

### Hyperparameters setting

For all cases, we train up to 1000 estimators at a learning rate of 0.05, with each tree allowed up to 31 leaves. To reduce overfitting, STAVAG randomly chooses 20% of genes for splitting and 90% of cells at each iteration.

### Significance of SVG, DVG, and TVG

The STAVAG score measures the overall contribution of each gene concerning spatial or temporal direction. To determine the significance of each gene, we employ a permutation-based approach. Suppose 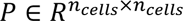 is a permutation matrix. We then have the permuted matrix:

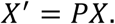

Here 𝑋^′^ has the same shape as 𝑋, but with rows rearranged. We next use 𝑋^′^ and 𝑌 as the input and calculate the aggregated STAVAG contribution score 𝑐𝑜𝑛𝑡𝑟𝑖𝑏𝑢𝑡𝑖𝑜𝑛^′^. At this point, 𝑋^′^ and 𝑌 are mismatched, resulting in the null distribution:

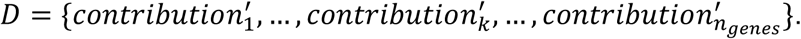

For gene 𝑘, its SVG significance score is defined as the top quantile within the null distribution 𝐷. Similarly, we can obtain the null distribution 𝐷_𝑗_ along the axis 𝑗 for DVG or TVG:

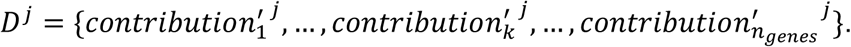

For gene 𝑘, its DVG/TVG p-value is defined as the proportion of values in the null distribution 𝐷_𝑗_ that are greater than or equal to the observed importance score. Mathematically, this can be expressed as:

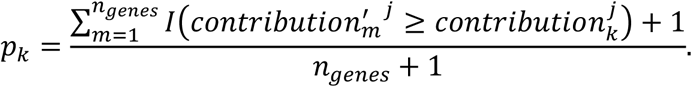

Here, 𝑝_𝑘_ is the p-value for the gene 𝑘, and 𝐼(⋅) is the indicator function, which equals 1 if the condition inside is true and 0 otherwise. In this study, we set 0.05 as the significance level. In our experiments, we found that this is a relatively stringent threshold. Users can adjust this value to obtain more significant variable genes as needed.

### Gene module detection

After identifying the DVG/TVG, we performed hierarchical clustering of the genes based on their pairwise correlations using the complete linkage method. The dendrogram’s distance is a user-defined parameter, which we set to 1 by default for global gene module detection. Users can decrease this parameter to obtain more refined gene modules. The expression of a given gene module is calculated as the average normalized gene expression of its involved genes.

### Enrichment analysis

We performed functional enrichment analysis with a web tool Metascape v3.5.20240901, which integrates multiple resources.

### Deconvolution of cSCC data

We collected the ST and scRNA-seq datasets of human cSCC from the original study^17^. We integrated these two datasets to estimate the proportion of various cell types in each spot of the ST dataset. Cell-type deconvolution of the ST dataset was performed using the R package RCTD. We set the parameters of the “create.RCTD” function as follows: max_cores = 10, UMI_min = 0, UMI_min_sigma = 0, MAX_MULTI_TYPES set to the number of cell types, CELL_MIN_INSTANCE = 0, and keep_reference = TRUE, while keeping all other parameters as default. For the “run.RCTD” function, we set doublet_mode = ‘full’ with all other parameters kept as default.

### Survival analysis

The TCGA HNSC samples were divided into high and low groups based on the threshold calculated using “surv_cutpoint” in the R package survminer V0.4.9. Kaplan-Meier survival curves were plotted using the “survfit” function.

### Variance method used in planarian data

We calculated a variance score for each gene based on section division of the anterior-posterior axis from the original study^2^. Let {𝑆_𝑛_}^10^ be the ten sections of the anterior-posterior axis divided by the original authors. The variance score for the gene 𝑘 was the variance of the average expression of each section:

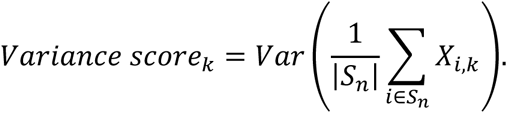

In this method, genes were ranked based on variance scores in descending order.

### Sample collection for acute myocardial infarction patients

We enrolled 88 patients who presented with chest pain within 3 hours of symptom onset. Subsequent coronary angiography confirmed acute myocardial infarction in 68 patients, while 20 patients were confirmed to have no severe coronary artery stenosis. The study was approved by the Ethics Committee of The First Affiliated Hospital of Ningbo University (No. 2021-R018-YJ01). The study was conducted in accordance with the principles outlined in the Declaration of Helsinki. All participants provided written informed consent for participation in the study.

### Proximity Extension Assay (PEA) Technology by Olink

Protein concentrations in plasma from 88 patients were measured using the Cardiometabolic and Cardiometabolic II panels (Olink Biosciences, Uppsala, Sweden), referring to a total of 729 proteins. The measurement was based on PEA technology, where plasma samples were incubated with antibodies conjugated with unique DNA oligonucleotide tags. When the antibodies bound to the target proteins, oligonucleotides were brought into proximity and extended by a DNA polymerase. The resulting DNA sequences were then quantified using the NovaSeq 6000 sequencing platform (Illumina, California, USA). Sequencing reads were normalized and presented as Normalized Protein eXpression (NPX) values.

### Code availability

The implementation of STAVAG can be accessed after publishing.

### Data availability

The data analyzed in this study can be accessed through: Human DLPFC 10x Visium dataset: http://spatial.libd.org/spatialLIBD; Human cSCC dataset: Gene Expression Omnibus (GEO) database with accession code GSE144240; Mouse STARmap cortex dataset and mouse MERFISH hypothalamic preoptic region dataset: https://www.bgiocean.com/vt3d_example/; Planarian 10x Visium 3D dataset: https://ngdc.cncb.ac.cn/STAPR/; Myocardial infarction 10x Visium dataset: GEO database with accession code GSE214611; Mouse placentation Stereo-seq datasets from E7.5 to E14.5: https://db.cngb.org/stomics/mpsta/; Mouse embryonic Stereo-seq datasets from E9.5 to E16.5: https://db.cngb.org/search/project/CNP0001543/.

