## Supplemental Text and Figures for "Uncovering directionally and temporally variable genes with STAVAG"

**Supplementary Notes**

### Supplementary Note 1: STAVAG demonstrates superior performance in identifying SVGs compared to other competing methods across 12 DPLFC datasets.

To benchmark STAVAG against other competing methods for the identification of SVGs, we first applied STAVAG to 12 human DLPFC 10x Visium datasets obtained from three donors[1]. These datasets were manually annotated with seven spatial domains including six cortical layers and white matter, arranged in a laminar organization (**Supplementary Fig.1a**).

To provide a fair evaluation metric, we took the intersection of the top 500 SVGs identified by each method. These genes are universally recognized as significant SVGs, thus their ranks within each method should be relatively high. We found that the ranks of these genes were the highest for STAVAG across the 12 datasets (median rank 95), demonstrating its ability to more effectively identify SVGs (**Supplementary Fig.1b, Supplementary Fig.2a**). Moreover, compared to the second-ranked method SPARK-X, STAVAG also showed a significant improvement (p-value = 1.6e-44). This superiority was consistently validated across all results from the top 300 to 700 SVGs, where STAVAG maintained a median rank from 65 to 140 (**Supplementary Fig.2b**). Further, leveraging the pre-annotated spatial domains, we calculated the spatial markers for each specific domain compared to others (**Supplementary Note 2, 3**). These markers, exhibiting clear spatial patterns, are expected to rank highly in the gene order produced by each method. In practice, these genes achieved the highest rankings in STAVAG (**Supplementary Fig.1c**). Under these two metrics, STAVAG and SPARK-X consistently performed at the first and second levels, while SpatialDE ranked third. Additionally, STAVAG and SPARK-X are the two fastest methods in terms of runtime (**Supplementary Fig.1d**).

### Supplementary Note 2: Definition of the spatial marker genes in DLPFC datasets.

Since the DLPFC dataset provides manually annotated spatial domains, we applied the Wilcoxon rank-sum test to identify the top 50 differentially expressed genes for each domain. The combined set of these genes was then used as the spatial marker genes for this dataset.

### Supplementary Note 3: Comparisons with other methods.

SPARK-X was applied using the R package SPARK v1.1.1 following its documentation at https://xzhoulab.github.io/SPARK/02_SPARK_Example/. HEARTSVG was applied using the R package HEARTSVG v1.1.0 following its documentation https://github.com/cz0316/HEARTSVG. SpatialDE was applied using the Python package SpatialDE v1.1.0 following its documentation https://github.com/Teichlab/SpatialDE. nnSVG was applied using the R package nnSVG v1.8.0 following its documentation at https://bioconductor.org/packages/release/bioc/vignettes/nnSVG/inst/doc/nnSVG.html.

**Supplementary Note 4: Comparison with SPARK-X on 3D ST data.**

We compared STAVAG with SPARK-X on the identification of SVGs, as SPARK-X is the only competing method that supports 3D ST data. We took the intersecting genes within the top 200–500 SVGs identified by both methods as representative SVGs and examined their ranks in each method. We found that these representative SVGs consistently ranked significantly higher in STAVAG than SPARK-X (p-values ranging from 8.2e-05 to 4.0e-19) (**Supplementary Fig.7b**). This indicates that STAVAG is more effective at accurately identifying SVGs in 3D spatial transcriptomics data.

**Supplementary Note 5: TVGs in the RZ region of MI data.**

We are also highly interested in the temporal gene expression changes within the RZ region, as it is present throughout all stages of disease progression. Using STAVAG, 121 TVGs were identified and subsequently grouped into three distinct modules (**Supplementary Fig. 10a, b**). RZ M1, enriched for mitochondrial genes, is predominantly expressed in the RZ at early times, reflecting its role in maintaining energy homeostasis in normal myocardium. RZ M2, enriched for ECM-related and cardiac remodeling genes, exhibits high expression at 72 and 168 hrs, highlighting its involvement in extracellular matrix remodeling and structural adaptation. RZ M3, enriched for cytoskeletal genes, heat shock proteins, and inflammatory factors, exhibits increased expression at 72 hrs. Indicating its role in cytoskeletal reorganization, stress response, and inflammation during tissue repair (**Supplementary Fig. 10d**). In short, STAVAG has provided a comprehensive understanding of the key functions associated with each stage of MI progression. These functions are consistent with the results of independent single-cell and spatial transcriptomic analyses[2, 3], further validating the stability and reliability of our method.

### Supplementary Figures

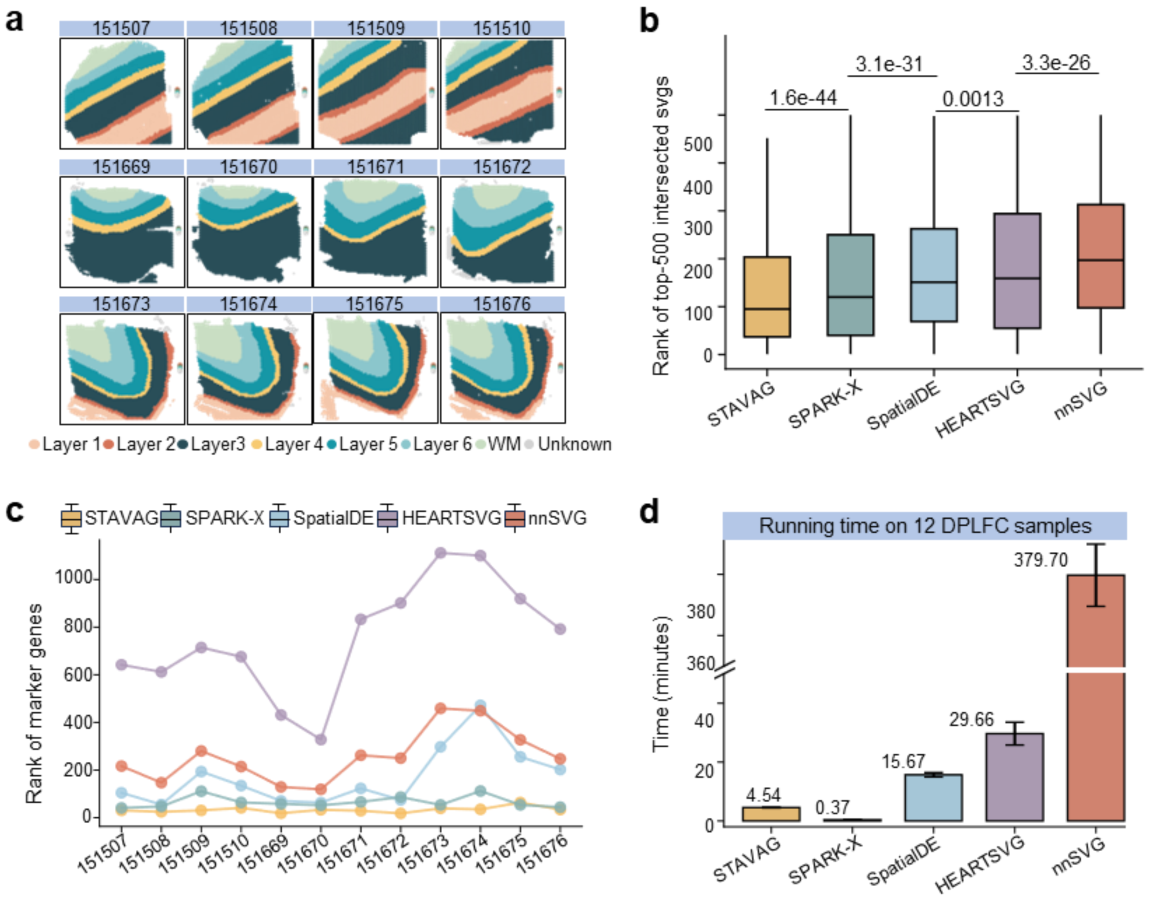


**Supplementary Figure 1. Benchmarking performance of STAVAG on identifying SVGs.** **a.** Overview of the 12 DLPFC datasets with its annotated spatial domains. **b.** Rank of the top-500 intersected SVGs for 12 DLPFC datasets, collected from STAVAG, SPARK-X, SpatialDE, HEARTSVG, nnSVG. **c.** Rank of the spatial marker genes calculated from the spatial domains. **d.** Running time of each method.

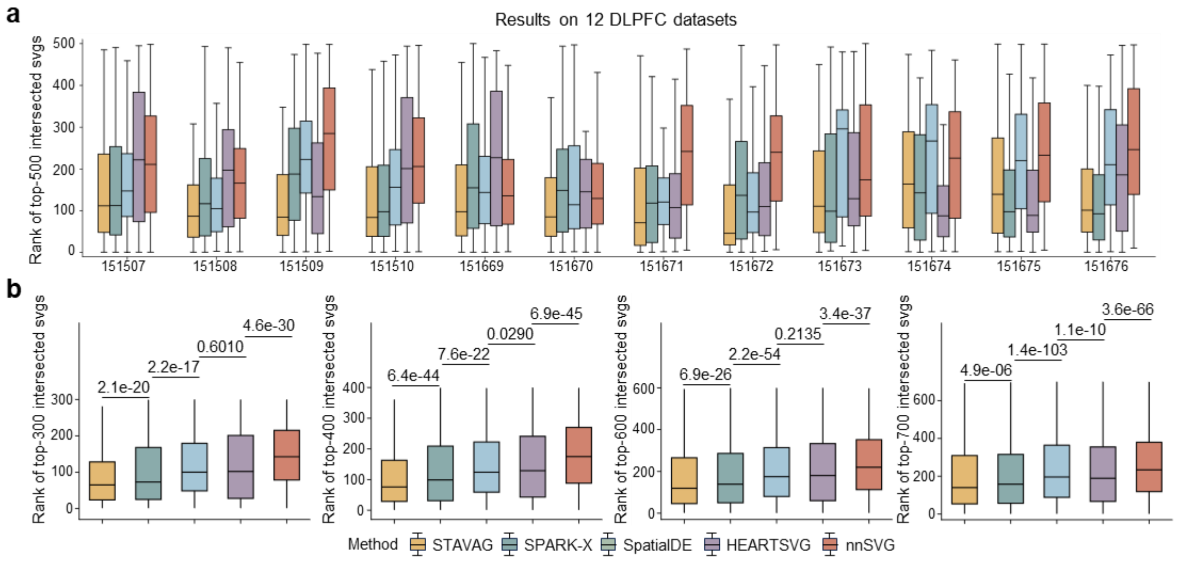


**Supplementary Figure 2. Evaluation on the performance of SVG detection for STAVAG and other competing methods. a.** Boxplots display the rank of the top-500 intersected SVGs for each DLPFC dataset, collected from STAVAG, SPARK-X, SpatialDE, HEARTSVG, nnSVG. **b.** Boxplots display the rank of the top-300, top-400, top-600, top-700 intersected SVGs for 12 DLPFC datasets, collected from STAVAG, SPARK-X, SpatialDE, HEARTSVG, nnSVG. For the boxplots in (**a**), and (**b**), center lines indicate median values and the lower and upper hinges represent 25th and 75th percentiles, respectively. The whiskers denote 1.5× the interquartile range.

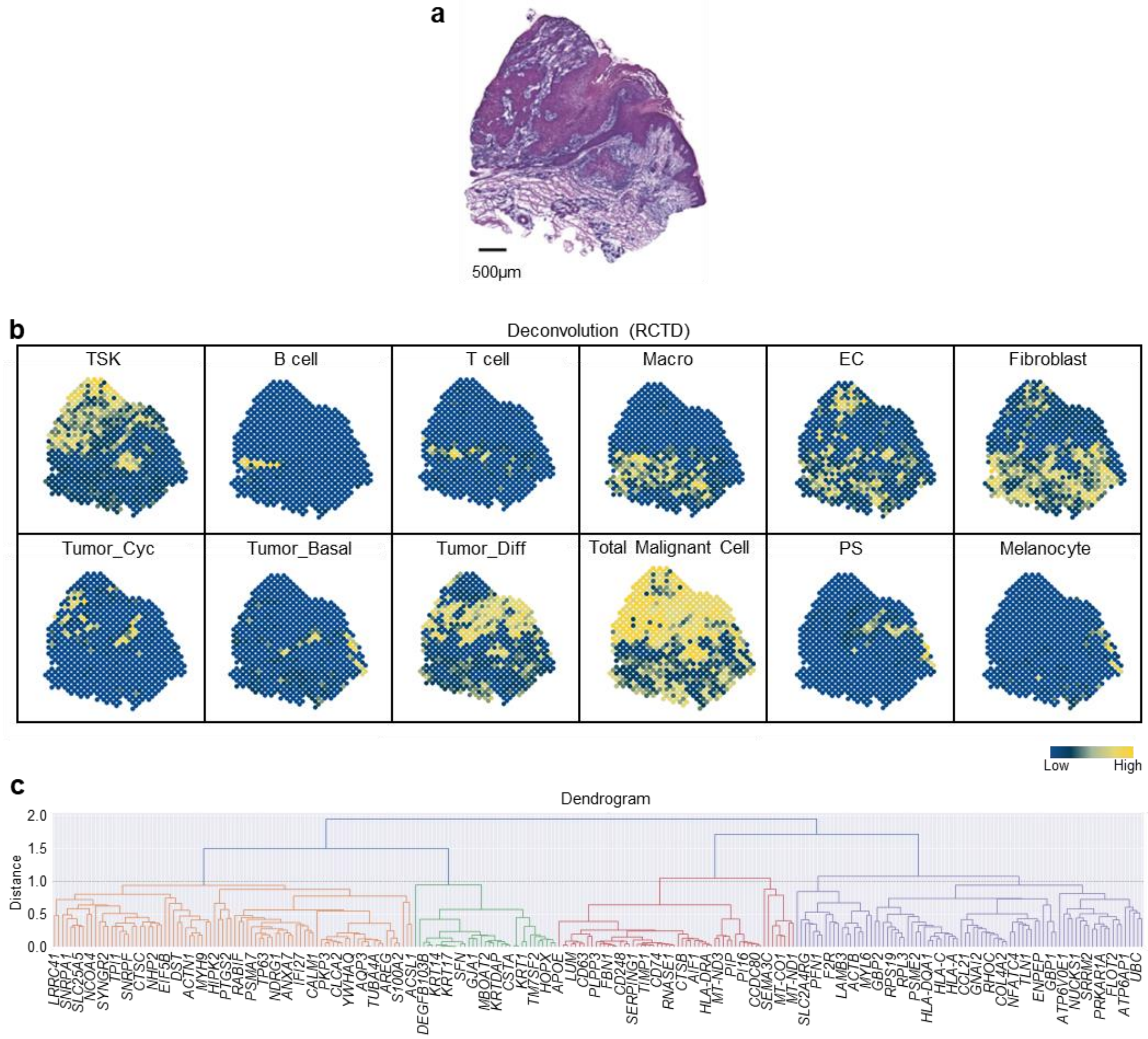


**Supplementary Figure 3. Evaluation on the DVG modules inferred from cSCC data. a.** Hematoxylin and eosin (H&E) staining of tissue section. **b.** The deconvolution results of cSCC data for different cell types, using RCTD. We use a dot plot to visualize the deconvolution probabilities. Here, Total Malignant Cell represents the aggregated deconvolution results of TSK, Tumor_Cyc, Tumor_Basal, and Tumor_Diff. **c.** Hierarchical clustering dendrogram showing the hierarchical relationships among the 266 DVGs along the y-axis of cSCC data. In the dendrogram, y-axis represents the distance between the clusters, measured by complete linkage.

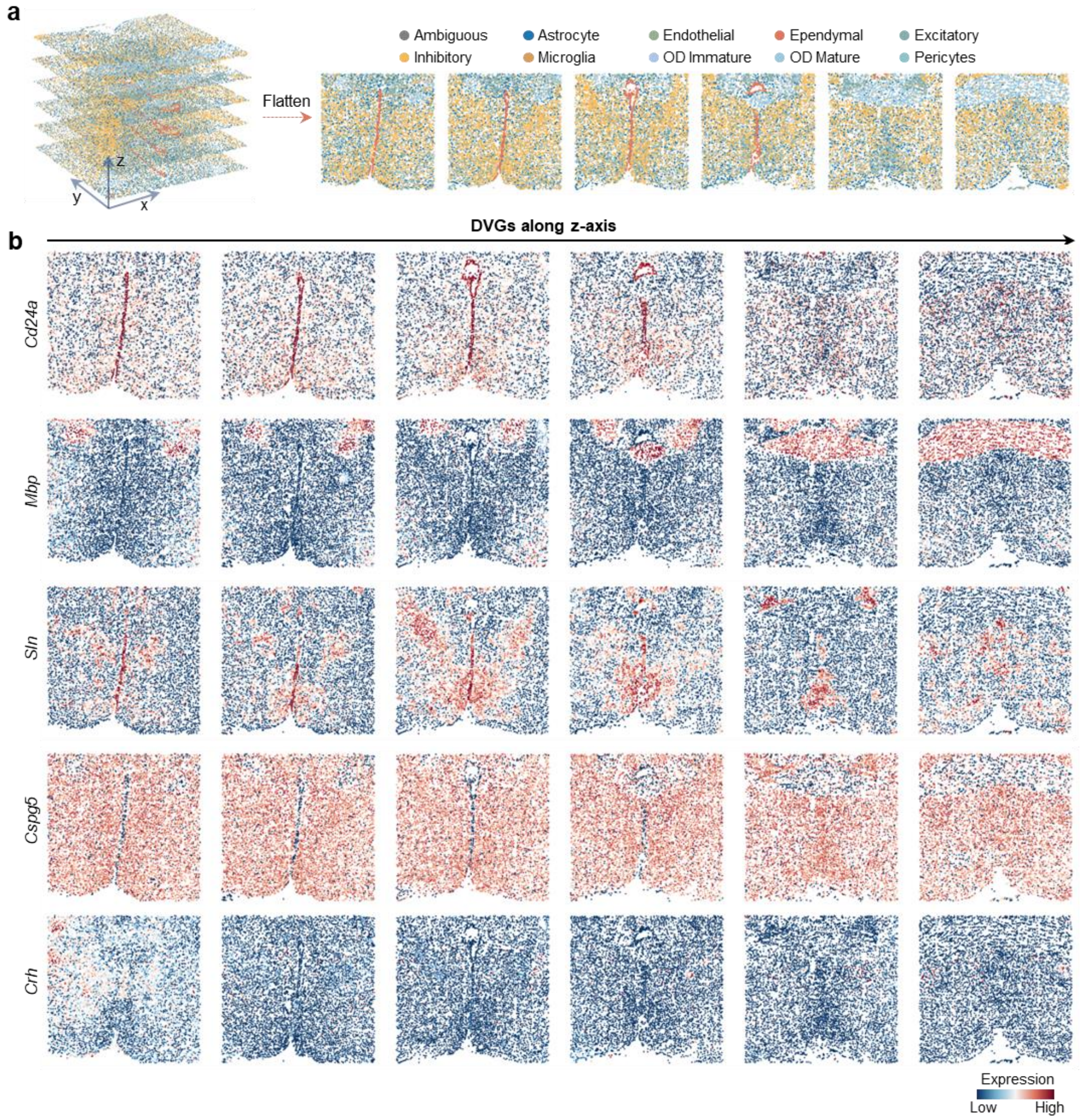


**Supplementary Figure 4. Evaluation on the representative DVGs inferred from MERFISH hypothalamic preoptic region data. a.** Overview of the MERFISH hypothalamic preoptic region data with its annotated cell types. To facilitate visualization, we selected slices 1, 3, 5, 7, 9, and 11 from the total 12 slices. **b.** The expression of representative DVGs along z-axis with different expression patterns.

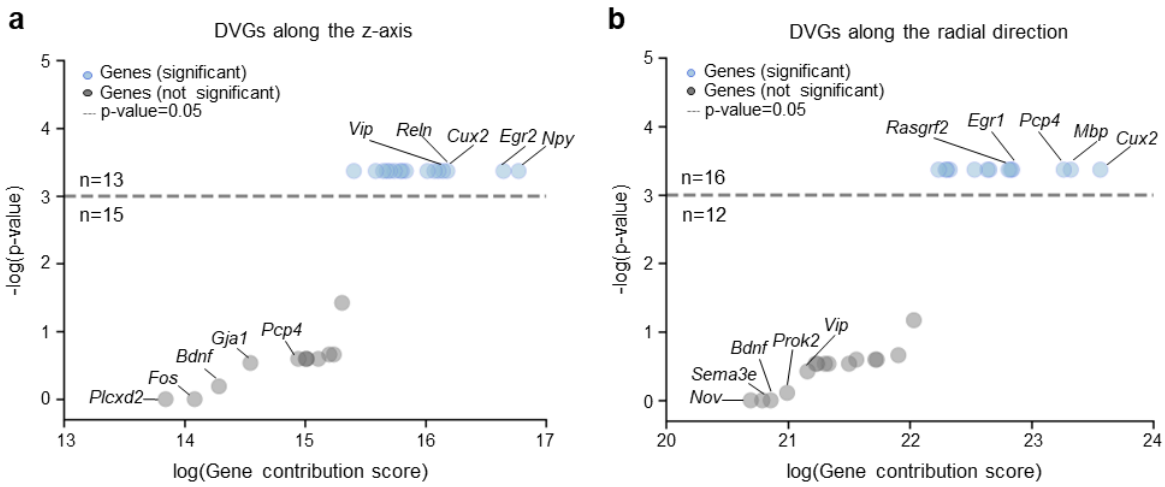


**Supplementary Figure 5. Representative DVGs from mouse cortex data. a-b.** Gene contribution score versus significance of STAVAG for all genes in the mouse cortex data. The STAVAG was applied to the z-axis (**a**) and the radial direction (**b**), respectively.

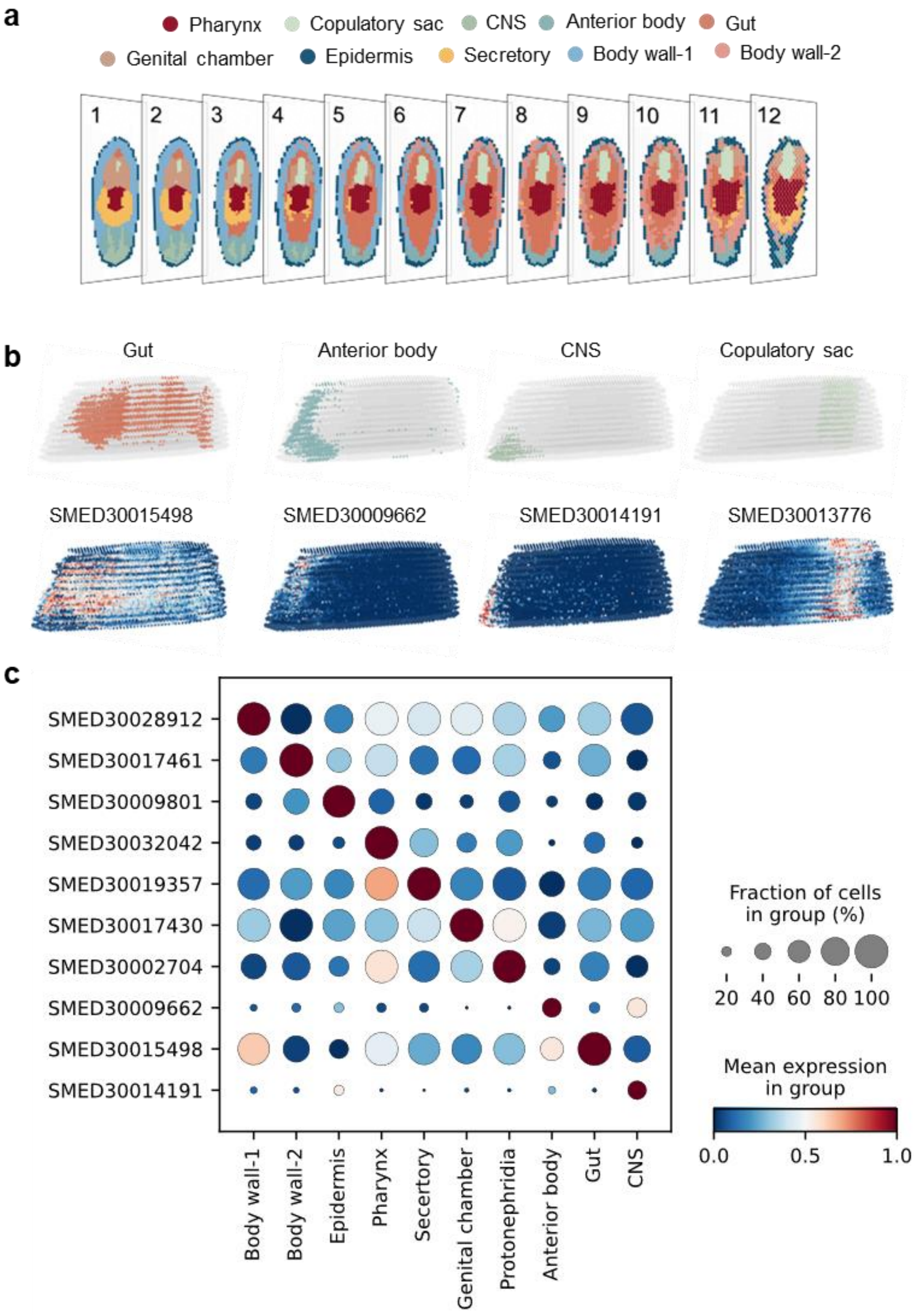


**Supplementary Figure 6. Representative DVGs from planarian data and their spatial distribution. a.** Sections 1 to 12 represent the directional transition from the ventral to the dorsal side. Each point represents a spot, with different colors indicating distinct domains. **b.** DVGs along DV axis with spatial distribution corresponding to Gut, Anterior body, CNS, and Copulatory sac. **c.** Dotplot displays the expressions of representative DVGs in **Fig. 4b** and **Supplementary Fig. 5a**.

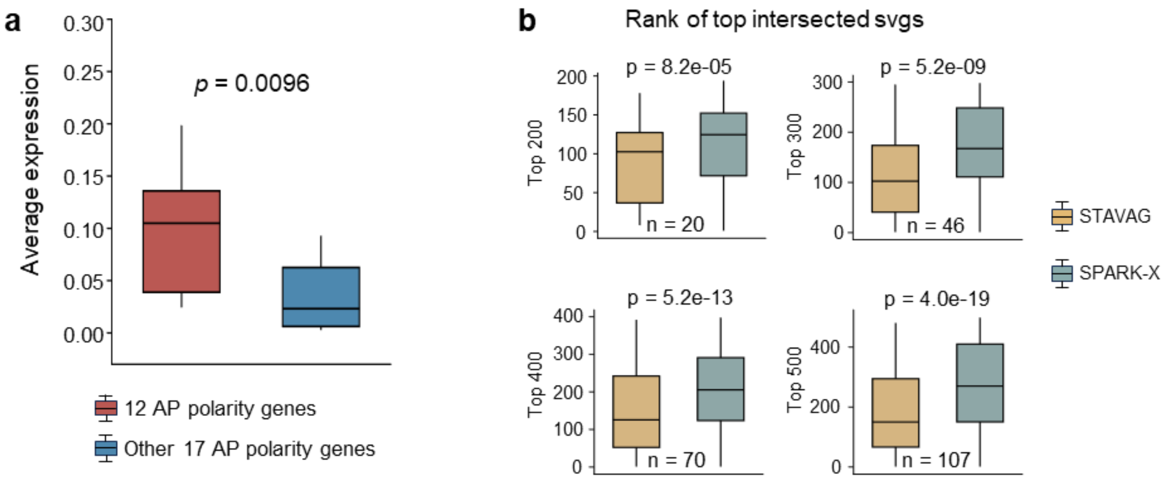


**Supplementary Figure 7. Evaluation on the DVGs and SVGs from planarian data. a.** Boxplot displays the expression of 29 AP polarity genes in spatial transcriptomics planarian data. These 29 AP polarity genes are divided into two groups: one consisting of 12 DVGs identified by STAVAG, and the other comprising the remaining genes. The p-value is calculated using a two-sided Wilcoxon rank-sum test. interquartile range. **b.** The boxplot illustrates the rankings of the top 200, 300, 400, and 500 intersecting SVGs identified by both STAVAG and SPARK-X in 3D planarian data. The p-value is calculated using a two-sided Wilcoxon rank-sum test. 'n' represents the number of intersecting genes identified as top SVGs by both methods. For the boxplot, center lines indicate median values, and the lower and upper hinges represent 25th and 75th percentiles, respectively. The whiskers denote 1.5× the interquartile range.

| SMED ID | Gene name | STAVAG_rank | SPARKX_rank | variance_rank |
|---|---|---|---|---|
| SMED30022285 | N/A | 9 | 1144 | 819 |
| SMED30011970 | N/A | 17 | 1148 | 396 |
| SMED30029487 | ndk | 20 | 2836 | 446 |
| SMED30000303 | Post-2c | 23 | 2763 | 2474 |
| SMED30028104 | ndl-2 | 40 | 2691 | 882 |
| SMED30010593 | wnt11-2 | 47 | 4029 | 4166 |
| SMED30006583 | fz4-1 | 50 | 4837 | 3409 |
| SMED30033839 | sp5 | 52 | 1918 | 1483 |
| SMED30015704 | Post-2d | 60 | 3364 | 2879 |
| SMED30031256 | wntP-2/wnt11-5 | 121 | 5113 | 1385 |
| SMED30025497 | sFRP-2 | 150 | 4568 | 1812 |
| SMED30020359 | ndl-3 | 217 | 3100 | 2624 |
| SMED30014748 | pkt7 | 1104 | 2016 | 762 |
| SMED30032613 | hox4b | 2046 | 5372 | 6294 |
| SMED30011055 | wntless | 3479 | 6080 | 5133 |
| SMED30002900 | N/A | 5704 | 7153 | 9425 |
| SMED30022468 | sFRP-1 | 5747 | 5062 | 3830 |
| SMED30033019 | ndl-5 | 9180 | 13389 | 13446 |
| SMED30032892 | foxD | 9370 | 8336 | 13430 |
| SMED30024252 | N/A | 10018 | 11567 | 15342 |
| SMED30026602 | wnt2 | 12196 | 9810 | 11440 |
| SMED30031321 | wntP-4/wnt11-3 | 12418 | 14170 | 14965 |
| SMED30007710 | wnt11-1 | 12653 | 10273 | 12546 |
| SMED30010051 | notum | 14257 | 10983 | 14777 |
| SMED30017913 | fz5/8-3 | 16557 | 7733 | 6112 |
| SMED30022929 | WntA | 17229 | 3415 | 1410 |
| SMED30021669 | Lox5a | 17693 | 11392 | 14589 |
| SMED30013810 | teashirt | 17929 | 7868 | 5535 |
| SMED30016952 | fz5/8-4 | 19152 | 4490 | 2694 |

**Supplementary Figure 8. The rank of the 29 AP polarity genes as determined by STAVAG, SPARK-X, and the Variance method.** Specifically, we compare the results of STAVAG with SPARK-X, which is the only competing method that supports 3D data. We also compare the results of STAVAG with Variance method (Methods), to see whether a simple variance-based method can effectively capture the AP polarity genes. The Gene name column indicates the symbols of homologous genes matched to other species by the sequence alignment tool BLAST, with N/A representing genes that cannot be matched.

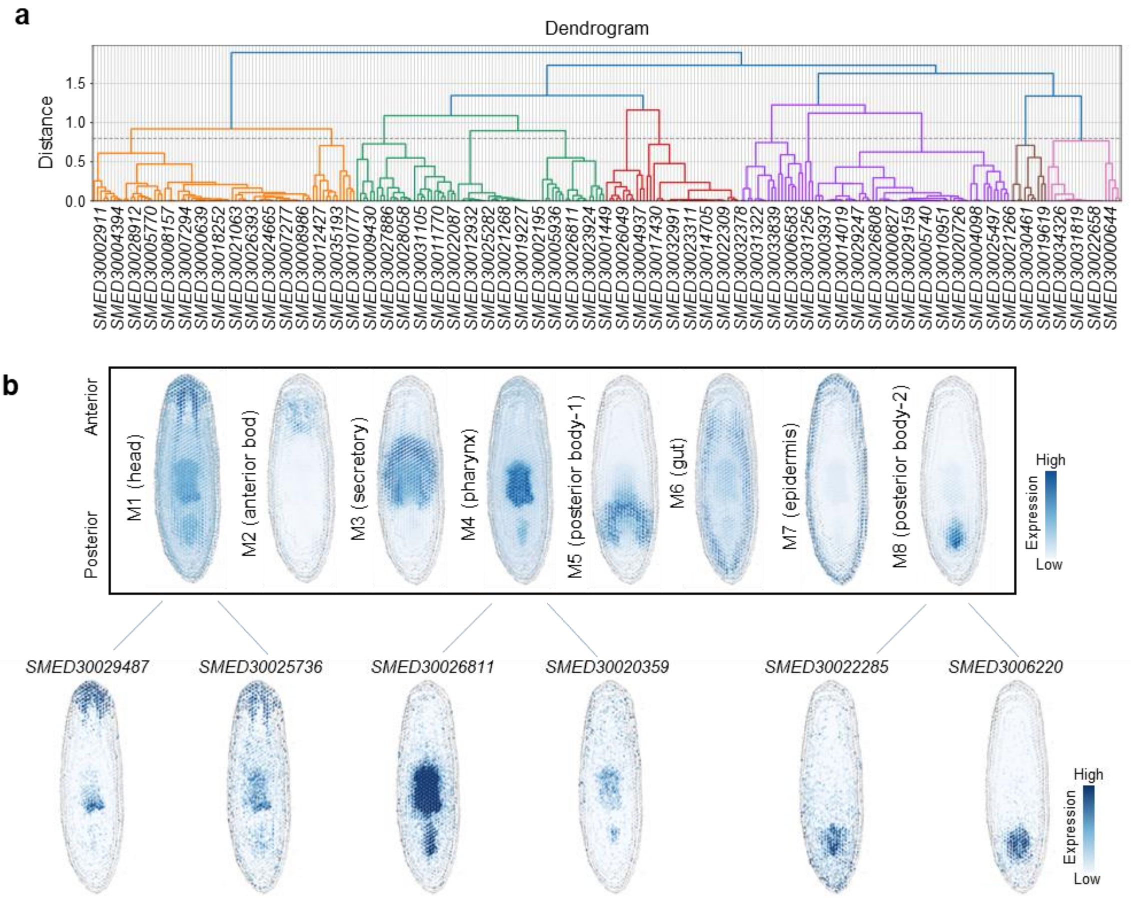


**Supplementary Figure 9. Evaluation on the representative DVG modules inferred from planarian data. a.** Hierarchical clustering dendrogram showing the hierarchical relationships among the DVGs identified from planarian data. In the dendrogram, y-axis represents the distance between the clusters, measured by complete linkage. **b.** Top-view visualization showing the expression of DVG modules and representative genes.

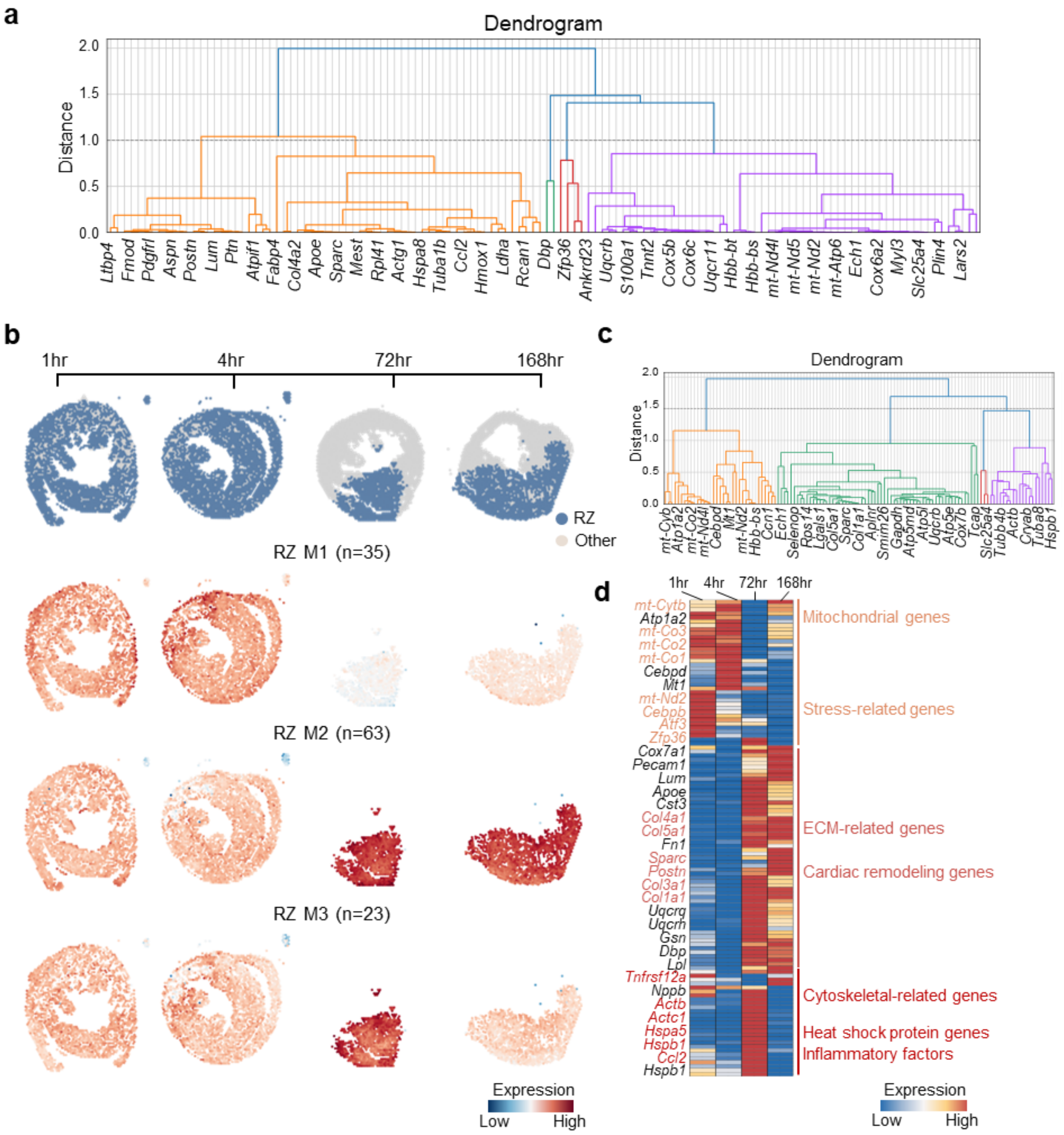


**Supplementary Figure 10. Evaluation on the representative TVG modules inferred from MI data. a.** Hierarchical clustering dendrogram showing the hierarchical relationships among the TVGs identified from MI data. In the dendrogram, y-axis represents the distance between the clusters, measured by complete linkage. **b.** Dotplot showing the expression of three TVG modules inferred from RZ region on spatial data. **c.** Hierarchical clustering dendrogram showing the hierarchical relationships among the 121 TVGs identified from RZ region in MI data. In the dendrogram, y-axis represents the distance between the clusters, measured by complete linkage. **d.** Heatmap showing the expression of TVGs identified from RZ region. Different colors were used to annotate gene-associated functions.

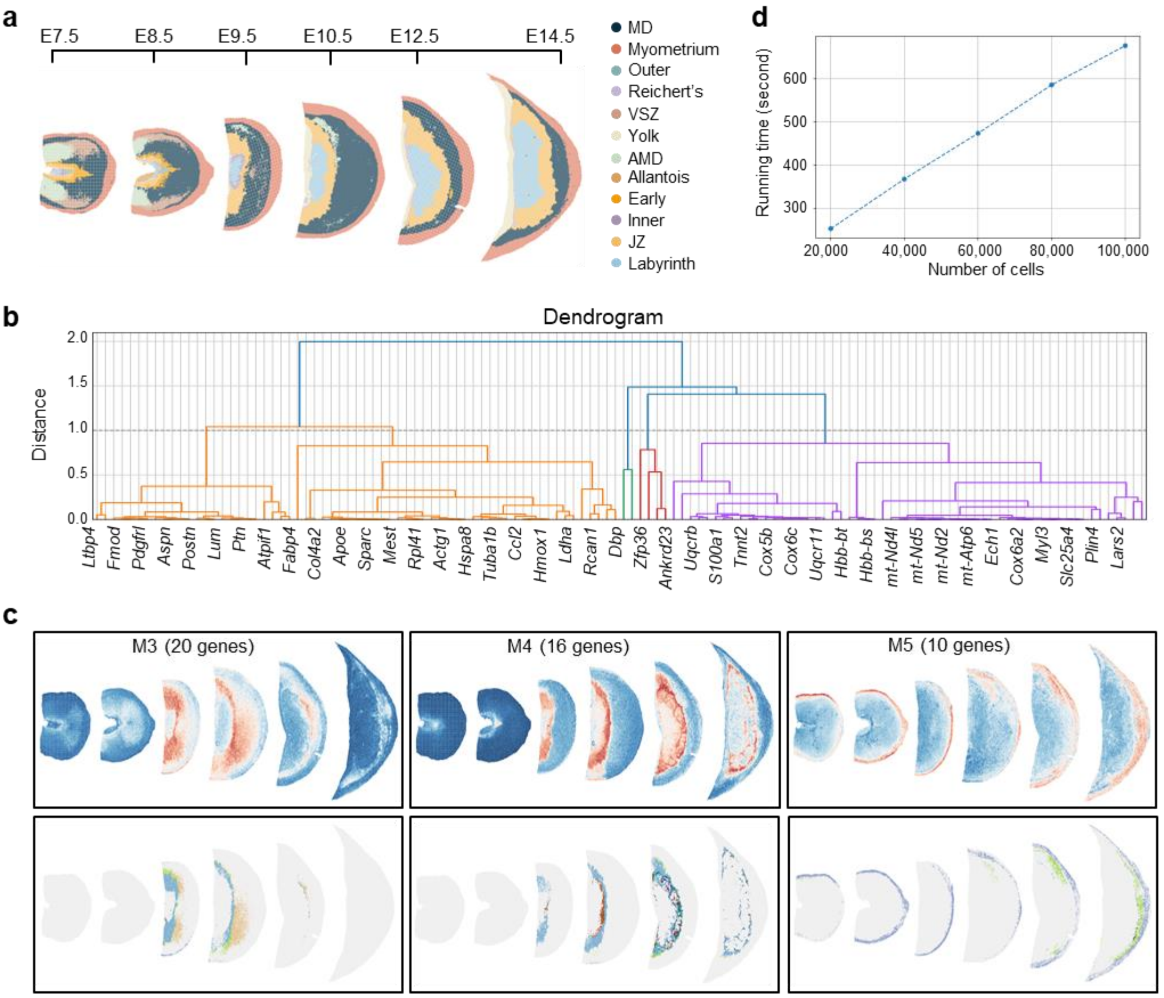


**Supplementary Figure 11. Evaluation on the representative TVG modules inferred from the mouse placentation development data and their functional analysis. a.** Scatterplot illustrating the spatial distribution of domains involved in mouse placentation. Each point represents a spot, with different colors indicating distinct domains. **b.** Hierarchical clustering dendrogram showing the hierarchical relationships among the 148 TVGs. **c.** Dotplot showing the expression of three TVG modules on spatial data (upper panel) and their primary expression regions (lower panel). **d.** Running time of STAVAG under different numbers of cells.

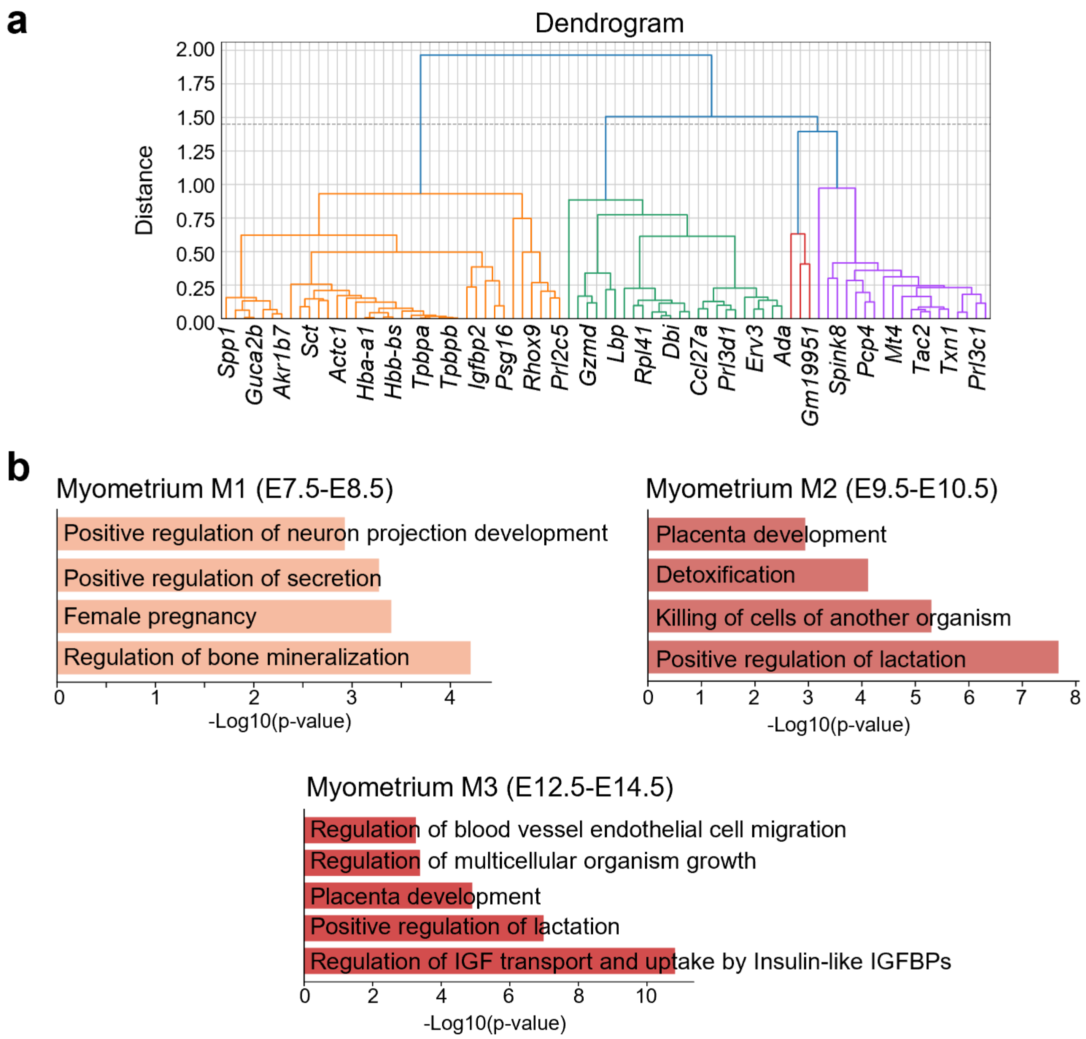


**Supplementary Figure 12. Evaluation on the representative TVG modules inferred from the myometrium of mouse placentation development data. a.** Hierarchical clustering dendrogram showing the hierarchical relationships among the 83 TVGs inferred from the myometrium region. **b.** Enrichment analysis of genes from Myometrium Module 1, 2, 3, presenting the top five terms for each module. IGF (Insulin-like Growth Factor), IGFBPs (Insulin-like Growth Factor Binding Proteins).

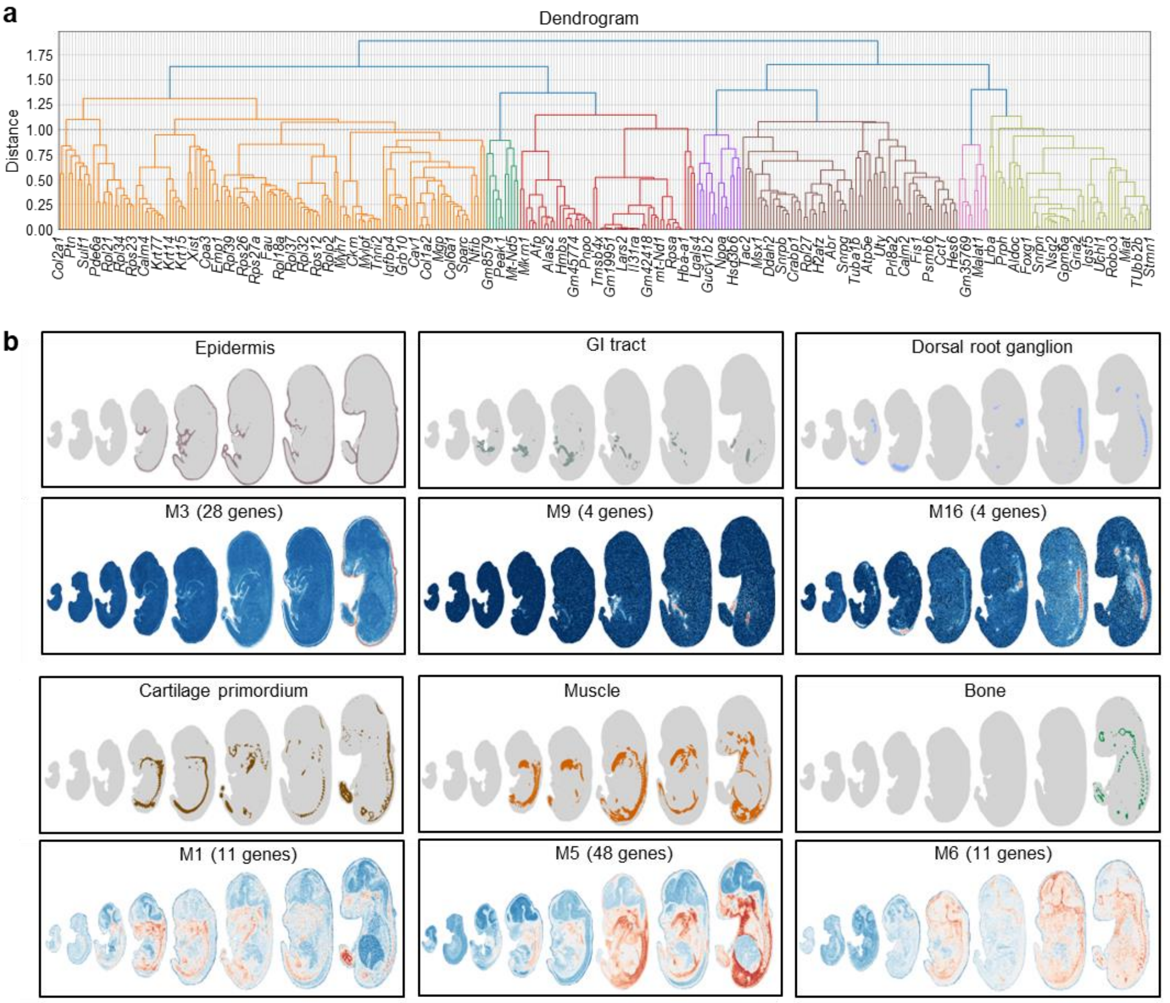


**Supplementary Figure 13. Evaluation on the tissue-specific TVG modules from mouse embryonic data. a.** Hierarchical clustering dendrogram showing the hierarchical relationships among the 358 TVGs. **b.** Dotplot showing the location of epidermis, GI tract, Dorsal root ganglion, Cartilage primordium, Muscle, and Bone, as well as the expression of TVG Module 3, 9, 16, 1, 5, 6 on spatial data.

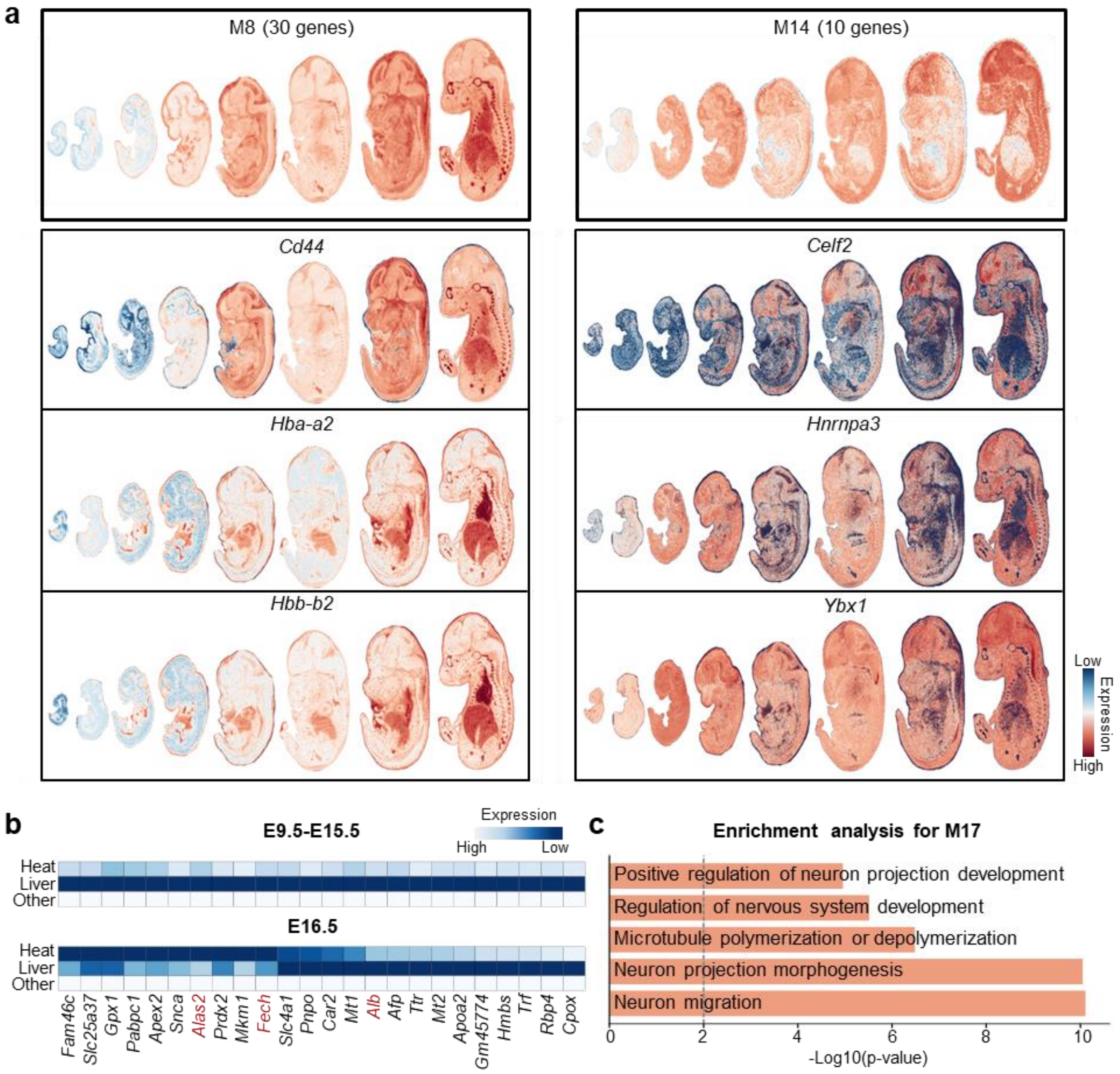


**Supplementary Figure 14. Evaluation on the global and tissue-interactive TVG modules from mouse embryonic data. a.** Dotplot showing the expression of TVG Module 8, 10, and representative genes on spatial data. **b.** Heatmap showing the expression of genes from M7 grouped by different tissues. Other means all tissues except the heart and liver. The genes highlighted in red represent those involved in heme biosynthesis. c. Enrichment analysis of genes from M17, presenting the top five terms.
